# The ER chaperone PfGRP170 is essential for asexual development and is linked to stress response in malaria parasites

**DOI:** 10.1101/406181

**Authors:** Heather M. Kudyba, David W. Cobb, Manuel A. Fierro, Anat Florentin, Dragan Ljolje, Balwan Singh, Naomi W. Lucchi, Vasant Muralidharan

**Author notes:** Address correspondence to Vasant Muralidharan.

## Abstract

The vast majority of malaria mortality is attributed to one parasite species: *Plasmodium falciparum*. Asexual replication of the parasite within the red blood cell is responsible for the pathology of the disease. In *Plasmodium*, the endoplasmic reticulum (ER) is a central hub for protein folding and trafficking as well as stress response pathways. In this study, we tested the role of an uncharacterized ER protein, PfGRP170, in regulating these key functions by generating conditional mutants. Our data show that PfGRP170 localizes to the ER and is essential for asexual growth, specifically required for proper development of schizonts. PfGRP170 is essential for surviving heat shock, suggesting a critical role in cellular stress response. The data demonstrate that PfGRP170 interacts with the *Plasmodium* orthologue of the ER chaperone, BiP. Finally, we found that loss of PfGRP170 function leads to the activation of the *Plasmodium* eIF2α kinase, PK4, suggesting a specific role for this protein in this parasite stress response pathway.

## INTRODUCTION

Malaria is a deadly parasitic disease that causes over 212 million cases and nearly 430,000 deaths each year, primarily in children under the age of five^1^. The deadliest human malaria parasite, *P. falciparum*, infects individuals inhabiting subtropical and tropical regions. These are some of the most impoverished regions of the world, making diagnosis and treatment challenging. Moreover, the parasite has evolved resistance to all clinically available drugs, highlighting an important need for uncovering proteins that are essential to the biology of this parasite^2–6^. Malaria is associated with a wide array of clinical symptoms, such as fever, chills, nausea, renal failure, pulmonary distress, cerebral malaria, and cardiac complications. It is the asexual replication of the parasite within the red blood cell (RBC) that is responsible for the pathology of the disease^7^.

In *P. falciparum*, the endoplasmic reticulum (ER) is a uniquely complex, poorly understood organelle. In fact, recent data suggest that ER proteins play a major role in resistance to the frontline antimalarial, artemisinin^8–10^. It is in this organelle that a variety of essential cellular functions occur, including protein trafficking, cellular signaling, and activation of stress response pathways^11–17^. Compared to other eukaryotes, the molecular mechanisms involved in these essential processes in *Plasmodium* remain poorly understood. Therefore, it is imperative to uncover proteins that regulate and maintain ER biology. One group of proteins likely governing many of these processes are ER chaperones^18–23^.Very little is known about the roles that ER chaperones play in *Plasmodium*, many of them defined merely based on sequence homology to other organisms. The *Plasmodium* genome encodes a relatively reduced repertoire of predicted ER chaperones, but it is predicted to contain two members of the conserved ER HSP70 chaperone complex, GRP78 (or BiP) and a putative HSP110 (PfGRP170 or PfHSP70-y)^24,25^. GRP170, in other eukaryotes, serves as nucleotide exchange factor for BiP^26,27^. Additionally, GRP170 has been reported to have holdase activity and can bind unfolded substrates independent of ATP or BiP^28–30^.

In this study, we used a conditional auto-inhibition strategy to generate conditional mutants for the putative ER chaperone, PfGRP170 (PF3D7_1344200)^31–33^. Using these conditional mutants, we localized PfGRP170 to the parasite ER, and show that unlike its orthologs in other eukaryotes, PfGRP170 is essential for parasite survival. Detailed life cycle analysis revealed that inhibition of PfGRP170 results in parasite death in early schizogony. The protein is required for surviving a brief heat shock, suggesting that PfGRP170 is essential during febrile episodes in the host. We show that despite a predicted transit peptide, PfGRP170 is not essential for protein trafficking to the apicoplast. Trafficking experiments using antibodies for two PEXEL Negative Exported Proteins and one protein containing a Plasmodium Export Element indicates that PfGRP170 is unlikely to be involved in protein export. Using a combination of mass spectroscopy approaches we identified potential interactors. Moreover, we demonstrate here that PfGRP170 interacts with the *Plasmodium* homolog of BiP (PF3D7_0917900) suggesting a conserved HSP70 ER chaperone complex. Finally, we show that conditional inhibition of PfGRP170 leads to the activation of the only known ER stress response pathway in Plasmodium, the PK4 pathway^10,16^.

## RESULTS

### PF3D7_1344200 is a putative GRP170 in P. falciparum

A blast search to identify ER localized Hsp70 proteins in *P. falciparum* revealed two proteins, HSP70-2 (PfGRP78/BiP) and a putative HSP110 (PF3D7_1344200). HSP110 proteins are considered large HSP70 chaperones, having sequence homology to both the nucleotide and substrate binding domains of other HSP70 members^34^. The increased size of HSP110 family members is the result of an extended α-helical domain at the C-terminus as well as an unstructured loop inserted in the substrate-binding domain^28,34^ (Figure 1A). In other eukaryotic organisms, the ER localized HSP110 (referred to as GRP170) is a chaperone with four primary protein domains: a signal peptide, a nucleotide binding domain, a substrate binding domain, and an extended C-terminus^26,34^. A protein sequence alignment using the yeast GRP170 (Lhs1) was used to predict the boundaries of these domains in PF3D7_1344200 (PfGRP170) (Figure 1A and Supplemental Figure 1). Most of the sequence conservation between Lhs1 and PfGRP170 was found to be in the nucleotide binding domain (Supplemental Figure 1). PfGRP170 is well conserved across multiple *Plasmodium* species, including other human malaria-causing species (Supplemental Figure 2). This level of conservation decreases in another apicomplexan (*T. gondii*) and even more so in yeast and humans (Supplemental Figure 2).

**Figure 1:**
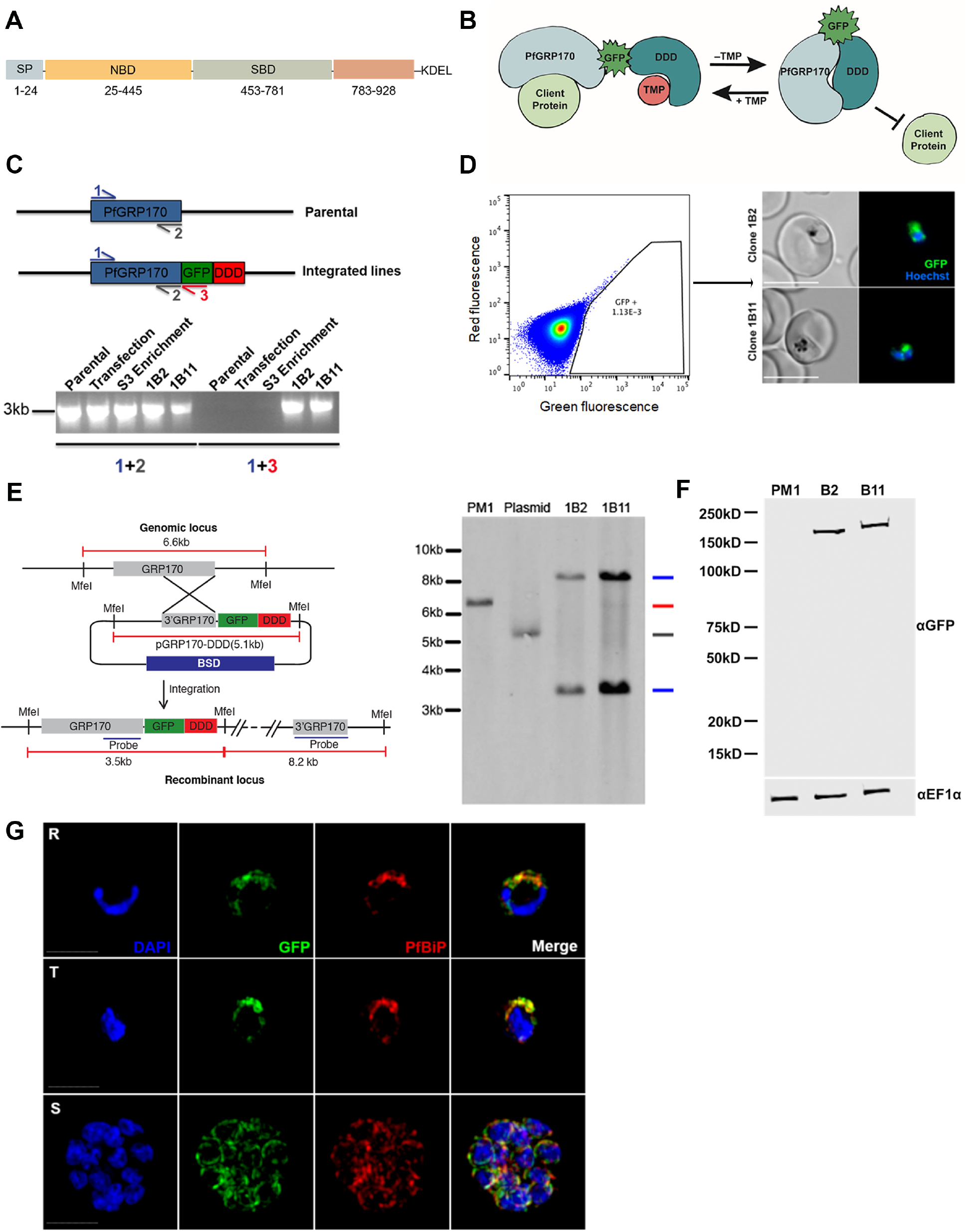
Generation of PfGRP170-GFP-DD Parasites. **(A)**. Schematic detailing the putative domain boundaries of PfGRP170 (PF3D7_1344200) based on the yeast homolog, Lhs1: Signal Peptide (SP), Nucleotide Binding Domain (NBD), Substrate Binding Domain (SBD), Extended C-terminus region (783-928), and an ER retention signal (KDEL). **(B)**. Schematic diagram demonstrating the conditional inhibition of PfGRP170. Conditional inhibition of PfGRP170 is achieved by the removal of Trimethoprim (TMP), which results in the unfolding of the destabilization (DDD). The chaperone recognizes and binds the unfolded DDD and is inhibited from interacting with client proteins. **(C)**. (Top) Schematic diagram of the PfGRP170 locus in the parental line (PM1KO) and the modified locus where PfGRP170 is endogenously tagged with GFP and DDD. Primers used for integration test and control PCR are indicated by arrows. The relative positions of Primer 1 (blue) and Primer 2 (Gray) on the PfGRP170 locus are shown. These two primers will amplify PfGRP170 in parental and transfected parasites. Primer 3 (Red) recognizes the GFP sequence. Primers 1 and 3 were used to screen for proper integration into the PfGRP170 locus. **(Bottom)** PCR integration test and control PCRs on gDNA isolated from the PM1KO (parental), the original transfection of the pPfGRP170-GFP-DDD plasmid after three rounds of Blasticidin (BSD) drug selection (Transfection), the PfGRP170-GFP-DDD transfected parasite lines after two rounds of enrichment for GFP positive cells (S3 enrichment), and PfGRP170-GFP-DDD clones 1B2 and 1B11 after MoFlo XDP flow sorting. The first 5 lanes are control PCRs using primers to amplify the PfGRP170 locus. The last 5 lanes are integration PCRs that only amplify if the GFP-DDD has been integrated into the genome. **(D). (Left)** MoFlo XDP flow data demonstrating the percentage of GFP positive parasites in transfected PfGRP170-GFP-DDD parasites following three rounds of drug selection with Blasticidin (BSD) and two rounds of enrichment with an S3 cell sorter. Using the MoFlo, single GFP positive cells were cloned into a 96 well plate. Two clones, 1B2 and 1B11, were isolated using this method. **(Right)** 1B2 and 1B11 parasites, were observed using live fluorescence microscopy. **(E).** Southern blot analysis of PfGRP170-GFP-DDD clones 1B2 and 1B11, PM1KO (parental control), and the PfGRP170-GDB plasmid is shown. Mfe1 restriction sites, the probe used to detect the DNA fragments, and the expected sizes are denoted in the schematic **(Left)**. Expected sizes for PfGRP170-GFP-DDD clones (blue), parental DNA (red), and plasmid (gray) were observed **(Right)**. Parental and plasmid bands were absent from the PfGRP170-GFP-DDD clonal cell lines. **(F)**. Western blot analysis of protein lysates from PM1KO (parental) and PfGRP170-GFP-DDD clonal cell lines 1B2 and 1B11 is shown. Lysates were probed with anti-GFP to visualize PfGRP170 and anti-PfEF1α as a loading control. **(G)**. Asynchronous PfGRP170-GFP-DDD parasites were paraformaldehyde fixed and stained with anti-GFP, anti-PfGRP78 (BiP), and DAPI to visualize the nucleus. Images were taken as a Z-stack using super resolution microscopy and SIM processing was performed on the Z-stacks. Images are displayed as a maximum intensity projection. The scale bar is 2μm.

### Generating PfGRP170-GFP-DDD conditional mutants

Conditional mutants for PfGRP170 (termed PfGRP170-GFP-DDD) were generated by tagging the endogenous PfGRP170 locus at the 3’ end, using single homologous crossover, with a GFP reporter, the *E. coli* DHFR destabilization domain (DDD), and an ER retention signal (SDEL) (Figure 1B). The endogenous PfGRP170 gene encodes a C-terminus SDEL sequence, a potential ER retention signal, and therefore we added an SDEL sequence after the DDD domain in order to avoid mislocalization of the tagged protein. In the presence of the small ligand Trimethoprim (TMP), the DDD is maintained in a folded state. However, if TMP is removed from the culture medium, the DDD unfolds and becomes unstable^31–33,35,36^. Intramolecular binding of the chaperone to the unfolded domain inhibits normal chaperone function (Figure 1B)^31–33^. Two independent transfections were carried out, and integrated parasites were selected via several rounds of drug cycling. PCR integration tests following drug selection indicated that the percentage of integrated parasites in both transfections were extremely low (Figure 1C). Consequently, standard limiting dilution could not be used to clone out integrated parasites. To circumvent this issue, flow cytometry was used to enrich and sort extremely rare GFP positive parasites. Despite low enrichments and sorting rates (GFP positive population =~1.13E-3), we successfully obtained two clones, termed 1B2 and 1B11, using flow sorting (Figure 1 C and D). Proper integration into the *pfgrp170* locus was confirmed by a Southern blot analysis (Figure 1E). Western blot analysis revealed that the PfGRP170-GFP-DDD protein was expressed at the expected size (Figure 1F). Immunofluorescence assays (IFA) and western blot analysis showed that the PfGRP170-GFP-DDD fusion protein was expressed and localized to the parasite ER during all stages of the asexual life cycle (Figure 1G and Supplemental Figure 3).

### PfGRP170 is essential for asexual growth and surviving febrile episodes

To investigate the essentiality of PfGRP170, PfGRP170-GFP-DDD asynchronous parasites were cultured in the absence of TMP, and parasitemia was observed using flow cytometry. A growth defect was seen within 24 hours after the removal of TMP, resulting in parasite death (Figure 2A). Furthermore, the two clonal parasite lines exhibited a dose-dependent growth response to TMP (Figure 2B). Consistent with data from other chaperones tagged with the DDD^31–33^, TMP removal did not result in degradation of PfGRP170-GFP-DDD (Figure 2C). Conditional inhibition of *Plasmodium* proteins that does not involve their degradation, has also been observed for other non-chaperone proteins ^37,38^. Moreover, the removal of TMP did not affect the ER localization of PfGRP170 (Supplemental Figure 4).

**Figure 2:**
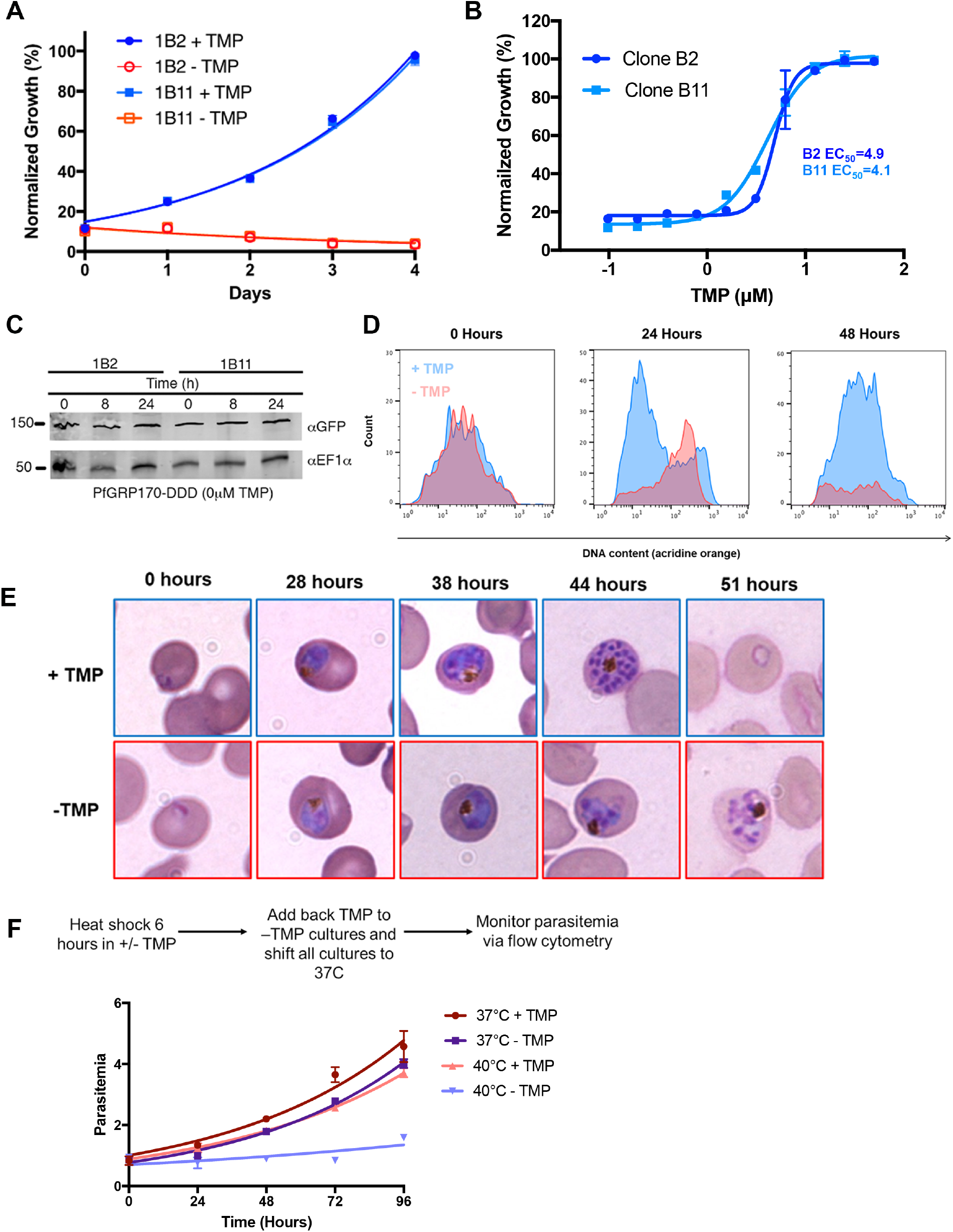
PfGRP170 is Essential and Required for Surviving a Heat Shock. **(A)**. Growth of asynchronous PfGRP170-GFP-DDD clonal cell lines 1B2 and 1B11, in the presence or absence of 20μM TMP, was observed using flow cytometry over 4 days. One hundred percent growth is defined as the highest parasitemia in samples with TMP, on the final day of the experiment. Data was fit to an exponential growth curve equation. Each data point is representative of the mean of 3 replicates ± S.E.M. **(B)**. Asynchronous PfGRP170-GFP-DDD clonal cell lines 1B2 and 1B11 were grown in a range of TMP concentrations for 48 hours. After 48 hours, parasitemia was observed using flow cytometry. One hundred percent growth is defined as the highest parasitemia in the presence of TMP on the final day of the experiment. Data was fit to a dose-response equation. Each data point is representative of the mean of 3 replicates ± S.E.M. **(C)**. Western blot analysis of PfGRP170-GFP-DDD lysates at 0, 8, and 24 hours following the removal of TMP is shown. Lysates were probed with anti-GFP to visualize PfGRP170 and anti-PfEF1α as a loading control. **(D)**. Flow cytometric analysis of asynchronous PfGRP170-GFP-DDD parasites, incubated with (Blue) and without TMP (Red), and stained with acridine orange. Data at 0, 24, and 48 hours after the removal of TMP are shown. **(E)**. TMP was removed from tightly synchronized PfGRP170-GFP-DDD ring stage parasites and their growth and development through the life cycle was monitored by Hema 3 stained thin blood smears. Representative images are shown from the parasite culture at the designated times. **(F)**. PfGRP170-GFP-DDD clones 1B2 and 1B11 were incubated with and without TMP for 6 hours at either 37°C or 40°C. Following the incubation, TMP was added back to all cultures and parasites were shifted back to 37°C. Parasitemia was then observed over 96 hours via flow cytometry. Data was fit to an exponential growth curve equation. Each data point shows the mean of 3 replicates ± S.E.M.

Using a nucleic acid stain, acridine orange, we used flow cytometry to specifically observe each stage of the asexual life cycle (ring, trophozoite, and schizont) in PfGRP170-GFP-DDD parasites incubated with and without TMP (Figure 2D). The amounts of RNA and DNA increase over the asexual life cycle as the parasite transitions from a ring to trophozoite to a multi-nucleated schizont. We observed that upon TMP removal, mutant parasites arrested in a relatively late developmental stage (Figure 2D). To identify the stage in the asexual life cycle where the mutant parasites died, TMP was removed from tightly synchronized ring stage cultures and parasite growth and morphology was assessed over the 48-hour life cycle. We observed morphologically abnormal parasites late in the lifecycle, when control parasites had undergone schizogony (Figure 2E). The PfGRP170-GFP-DDD parasites grown without TMP ultimately failed to progress through schizogony and did not reinvade new RBCs (Figure 2E).

The cytoplasmic ortholog of PfGRP170, PfHSP110, was previously shown to be essential for surviving heat stress^33^. Therefore, we tested whether PfGRP170 mutants were sensitive to a brief heat shock. Asynchronous parasites were incubated in the absence of TMP for 6 hours at either 37°C or 40°C. Following the 6-hour incubation, TMP was added back to all cultures, which were then grown at 37°C for two growth cycles, while measuring parasitemia every 24 hours. The growth of parasites at 37°C was not significantly affected by the brief removal of TMP (Figure 2F). In contrast, incubating parasites at 40°C without TMP resulted in reduced parasite viability compared to parasites grown at 40°C with TMP (Figure 2F).

### PfGRP170 is not required for trafficking of apicoplast proteins

Protein trafficking to the apicoplast is essential for parasite survival. Proteins targeted to the apicoplast contain an N-terminal transit peptide that is revealed upon signal peptide cleavage in the ER^15,39^. It remains unclear whether apicoplast targeted proteins go through the Golgi before reaching their final destination. It has been shown that disruption of ER to Golgi trafficking, using Brefeldin A (BFA), does not reduce apicoplast transport^15,40^. One of these studies further demonstrated that the addition of an ER retention sequence (SDEL), to a GFP with a transit peptide, did not reduce apicoplast trafficking or transit peptide cleavage^40^. However, a separate but similar analysis came to the opposite conclusion^41^. Thus, the identification, packaging, and transport of apicoplast-targeted proteins from the ER remain unanswered questions.

Two software analysis tools (Prediction of Apicoplast-Targeted Sequences (PATS) and PlasmoAP) predicted a strong apicoplast transit peptide for PfGRP170, despite our observation of a definite ER localization (Figure 1G and Figure 3A). We were therefore interested to find out whether PfGRP170 plays a role in apicoplast trafficking. We tested whether the putative PfGRP170 transit peptide (amino acids: 1 to 150) could be trafficked to the apicoplast by episomally expressing the predicted PfGRP170-transit peptide fused to a GFP reporter without an ER retention signal (Figure 3B). We performed co-localization assays using ER, apicoplast, and Golgi markers and determined that the putative transit peptide localized to the ER due to colocalization with ER marker Plasmepsin V (Figure 3B).

**Figure 3:**
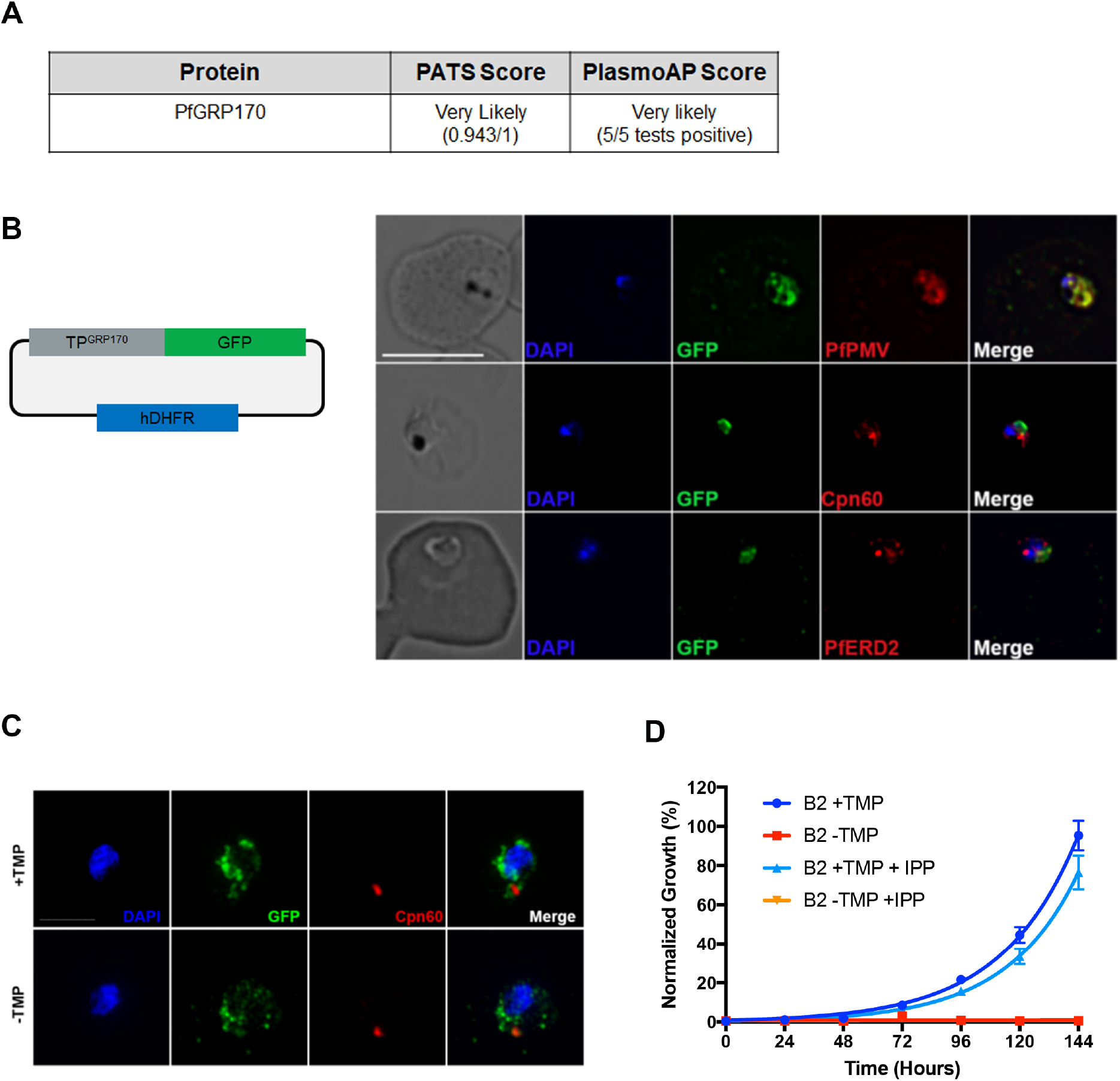
Putative PfGRP170 apicoplast transit peptide localizes to the ER and conditional inhibition of PfGRP170 does not affect trafficking of apicoplast proteins. **(A)**. Analysis of PfGRP170’s protein sequence using two apicoplast transit peptide prediction programs: Prediction of Apicoplast-Targeted Sequences (PATS) and PlasmoAP. **(B)**. PfGRP170’s putative apicoplast transit peptide was fused to GFP and transfected into 3D7 parasites. Parasites were fixed with acetone and stained with DAPI, anti-GFP (to label the PfGRP170 putative transit peptide) and either anti-PfPMV (ER), anti-PfERD2 (Golgi), or anti-Cpn60 (Apicoplast) to determine subcellular localization. The images were taken with Delta Vision II, deconvolved, and are displayed as a maximum intensity projection. The scale bar is 5μm. **(C)**. Synchronized ring stage PfGRP170 parasites were incubated for 24 hours with and without TMP. Following the incubation, the parasites were fixed with paraformaldehyde and stained with DAPI, anti-GFP (PfGRP170) and anti-Cpn60 (Apicoplast). Images were taken as a Z-stack using super resolution microscopy and SIM processing was performed on the Z-stacks. Images are displayed as a maximum intensity projection. The scale bar is 2μm. **(D)**. Asynchronous PfGRP170-GFP-DDD parasites were incubated with and without TMP and in the presence or absence of 200μM IPP. Parasitemia was monitored using flow cytometry for 144 hours. One hundred percent growth is defined as the highest parasitemia in the presence of TMP, on the final day of the experiment. Data was fit to an exponential growth curve equation. Each data point is representative of the mean of 3 replicates ± S.E.M.

To determine the role of PfGRP170 in trafficking proteins to the apicoplast, we removed TMP from PfGRP170-GFP-DDD parasites and examined the localization of the apicoplast-localized cpn60^32,42,43^. No defects in apicoplast localization of cpn60 were observed (Figure 3C). Additionally, incubation with the essential apicoplast metabolite IPP^44^, failed to rescue or have any positive effect on TMP removal in PfGRP170-GFP-DDD parasites (Figure 3D).

### Interactions of PfGRP170

Two independent approaches were taken to identify the interacting proteins of PfGRP170 (Figure 4A). The first was an anti-GFP Immunoprecipitation (IP) followed by mass spectroscopy. In the second approach we generated a parasite line episomally expressing PfGRP170 tagged with an HA, the promiscuous Biotin Ligase (BirA), and an ER retention signal (KDEL). When exogenous biotin is added to the PfGRP170-HA-BirA parasites, the BirA tagged protein will biotinylate interacting proteins or those that are in close proximity^45^. These biotinylated proteins were isolated using streptavidin coated magnetic beads. A western blot analysis confirmed expression of the PfGRP170-HA-BirA fusion protein (Supplemental Figure 5A). Colocalization IFA’s confirmed that the PfGRP170-HA-BirA protein localizes to the parasite ER (Supplemental Figure 5B). Additionally, western blot analysis demonstrates that proteins are biotinylated in the PfGRP170-HA-BirA parasite lines when biotin is added (Supplemental Figure 5C).

**Figure 4:**
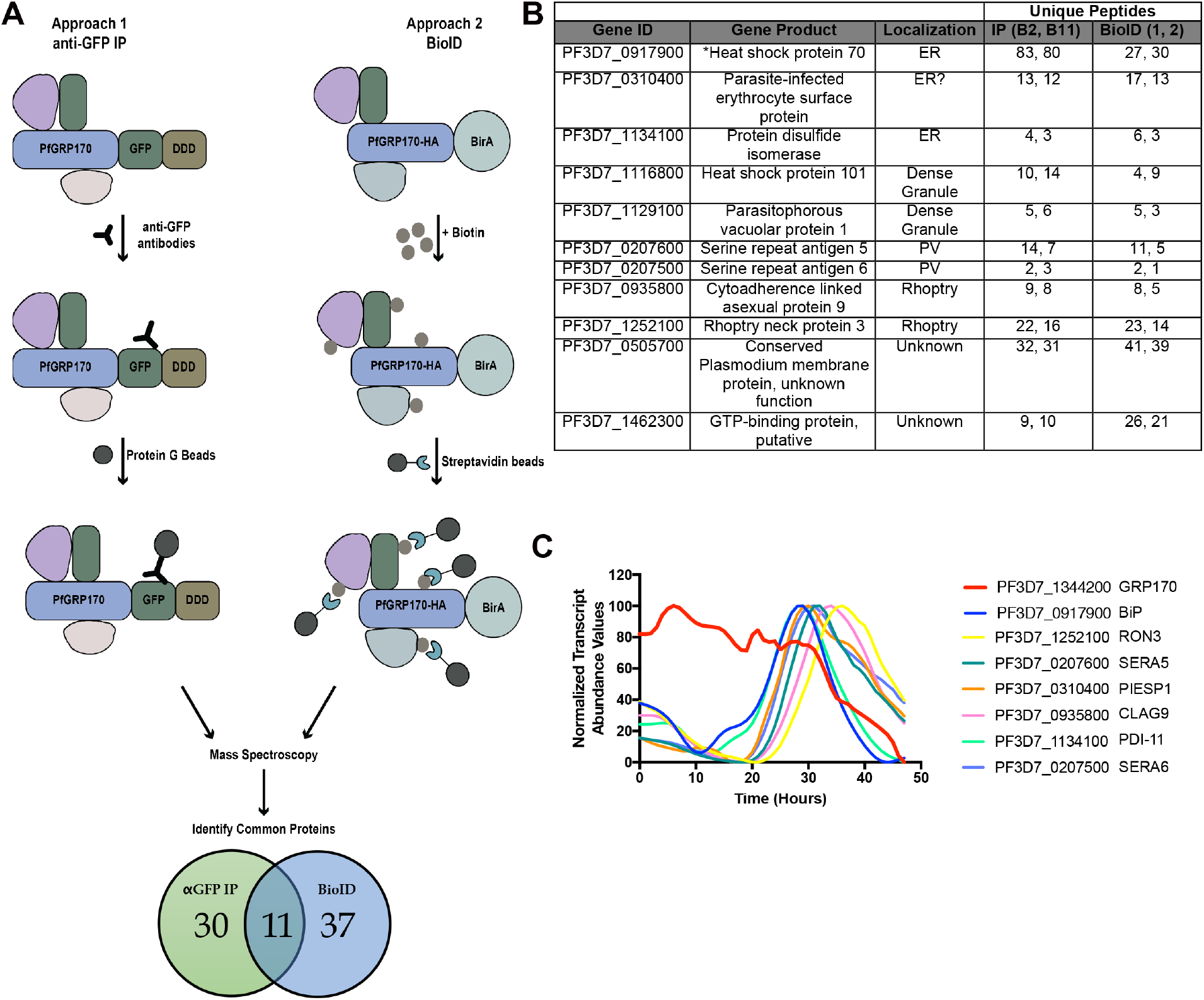
PfGRP170 interacting partners. **(A)**. Schematic diagram illustrating the two independent methods used to identify potential interacting partners of PfGRP170: anti-GFP Immunoprecipitation (IP) using lysates from PfGRP170-GFP-DDD parasites and streptavidin IP of PfGRP170-BirA parasites incubated with biotin for 24 hours followed by mass spectroscopy. The proteins identified from each IP were filtered to include only proteins that had a signal peptide and/or transmembrane domain using PlasmoDB. Proteins found in the respective control IP’s (excluding PfBiP) were also removed from further analysis (data in Supplemental Table 1). **(B)**. The 11 proteins identified in both independent mass spectroscopy approaches (See Figure 4A and Supplemental Table 1). The PlasmoDB gene ID, gene product, putative subcellular localization, and number of unique peptides identified for each protein in each independent experiment are listed. **(C)**. The relative transcript abundance of interacting proteins, with peak expression around the time the PfGRP170-GFP-DDD parasites die (36-44 hours), are plotted using genome-wide real-time transcript data^46^.

Mass spectroscopy was used to identify PfGRP170 interacting proteins from two independent anti-GFP IP’s of PfGRP170-GFP-DDD parasites and two biological replicates of streptavidin pulldowns from PfGRP170-HA-BirA parasites following an incubation with biotin (Supplemental Table 1). Further, two independent anti-GFP IP’s from the PM1 parental control and one streptavidin pull down from 3D7 parasites incubated with biotin to filter out non-specific interactions (Supplemental Table 1). Both BiP and PfGRP170 were found in the control IP’s, albeit in lower peptide counts (Supplemental Table 1). This is not surprising as chaperones are common contaminants in mass spectroscopy. However, due to the documentation of PfGRP170 being an interactor and regulator of BiP function, we opted to keep BiP in our analysis^26^. In order to obtain a list of proteins specific to the ER and parasite secretory pathway, proteins identified by mass spectroscopy in each IP were filtered to include only those, which had a signal peptide or transmembrane domain. Thirty proteins were found in both the 1B2 and 1B11 in the anti-GFP IP’s and 37 proteins were found in the two replicates of the PfGRP170-HA-BirA streptavidin pull down (Figure 4A). Of these, 11 proteins were identified using both approaches suggesting that these are true interactors of PfGRP170 (Figure 4B). Using recently published real-time transcriptional abundance data, we plotted the normalized transcriptional abundance values for all 11 proteins^46^ (Supplemental Figure 6). Upon removal TMP, the PfGRP170-DDD parasites die 38-44 hours post invasion (Figure 2E) and therefore, it is likely that the essential function of PfGRP170 is linked to proteins expressed during these late stages of the asexual life-cycle. Excluding proteins which were expressed earlier in the life cycle, narrows the list of putative essential interactors of PfGRP170 to the seven proteins (Figure 4C)^46^.

### PfGRP170 is not required for trafficking to the RBC

In order for the parasite to grow, develop, and divide, it must drastically remodel the host RBC^12^. These modifications are accomplished through the export of proteins from the ER to the RBC. In model eukaryotes, such as yeast and mammalian cells, molecular chaperones, and specifically those that are ER-localized, play central roles in protein trafficking^18,19^. Therefore, we tested whether conditional inhibition of PfGRP170 would prevent trafficking of several exported proteins (PfHSP70x, PfMAHRP1, and FIKK4.2). Our results demonstrate that loss of PfGRP170 function did not affect the localization of these proteins to the host RBC (Supplemental Figure 7A-C).

### PfGRP170 and BiP interact

One of the most abundant proteins identified in our mass spectroscopy data was PfBiP (Figure 4B and Supplemental Table 1). However, PfBiP was also found in our IPs performed with parental controls, therefore, we were tested whether PfGRP170 and PfBiP interact in *P. falciparum*. We performed an anti-GFP Co-IP and probed the lysate for PfBiP. We observed that PfGRP170 and PfBiP interact and this interaction is not lost upon TMP removal (Figure 5A). As a control, we probed the GFP Co-IP lysates for a different ER protein, Plasmepsin V (PMV), and found that it did not pull down with PfGRP170 (Figure 5B).

**Figure 5:**
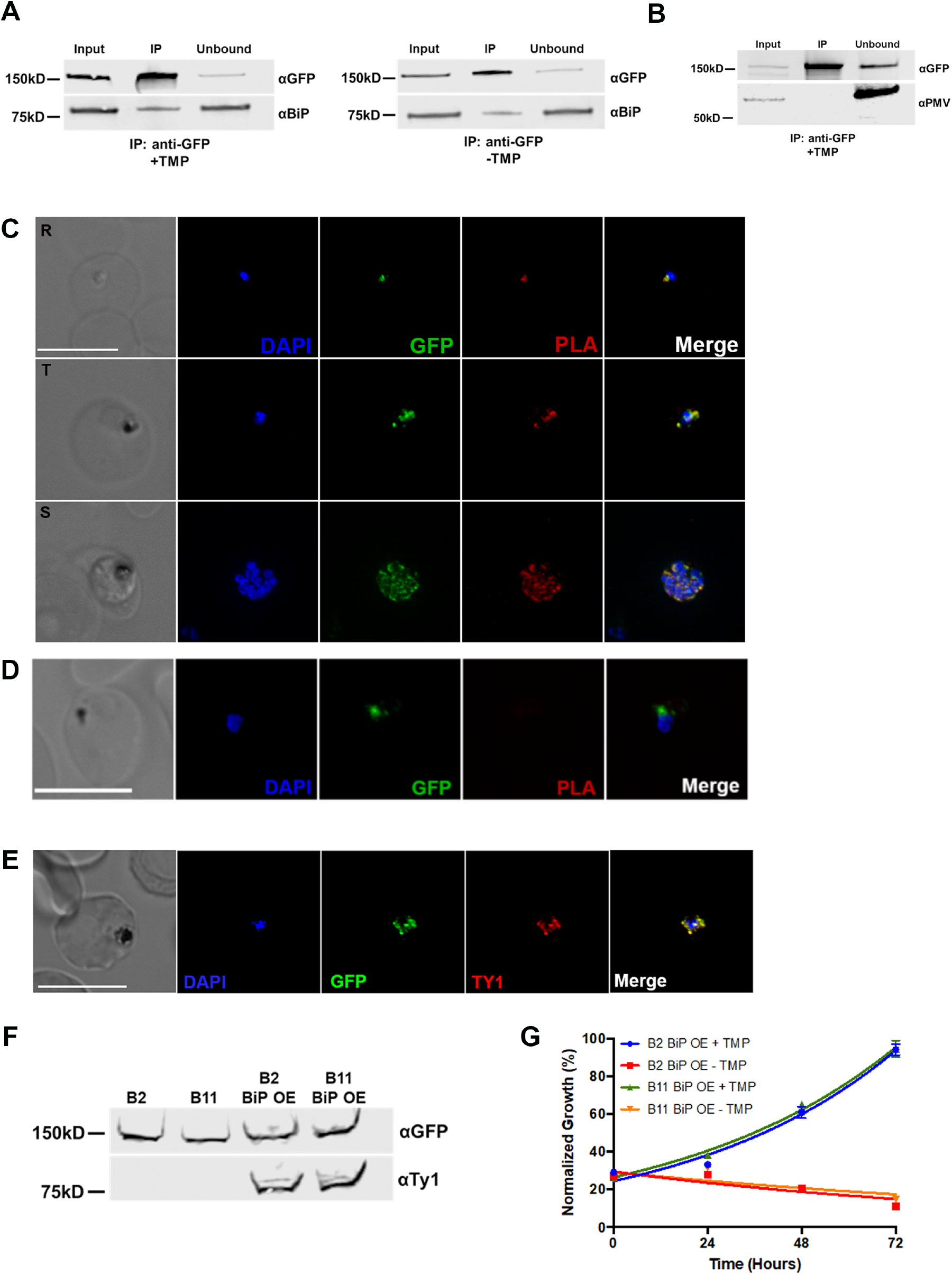
PfGRP170 Interacts with BiP. **(A)** Synchronized ring stage PfGRP170-GFP-DDD parasites were incubated with and without TMP for 24 hours. Following this incubation, an anti-GFP IP was performed and input, IP, and unbound fractions were analyzed using a western blot. The blot was probed using anti-GFP and anti-BiP. **(B).** Western blot analysis of an anti-GFP IP performed on asynchronous PfGRP170-GFP-DDD parasites. Input, IP, and unbound fractions are shown. The blot was probed using anti-GFP and anti-PfPMV. **(C).** *In vivo* interaction of PfGRP170 and BiP. PfGRP170-GFP-DDD parasites were paraformaldehyde fixed and stained with anti-GFP and anti-BiP. A Proximity Ligation Assay (PLA) was then performed. The scale bar is 5μm. A negative control using anti-GFP and anti-PfPMV is shown in **(D).** **(E).** Asynchronous PfGRP170-GFP-DDD parasites overexpressing PfBiP-Ty1were paraformaldehyde fixed and stained with anti-GFP (PfGRP170), anti-Ty1 (PfBiP-Ty1-KDEL), and DAPI to visualize the nucleus. The images were taken with Delta Vision II, deconvolved, and are displayed as a maximum intensity projection. The scale bar is 5μm. **(F).** Western blot analysis of protein lysates from parental 1B2 and 1B11 parasites as well as 1B2 and 1B11 parasites overexpressing the PfBiP-Ty1fusion protein. Lysates were probed with anti-GFP to visualize PfGRP170 and anti-Ty1 to visualize PfBiP-Ty1-KDEL. **(G).** Parasitemia of asynchronous PfGRP170-GFP-DDD parasites expressing PfBiP-Ty1-KDEL, in the presence or absence of 20μM TMP, was observed using flow cytometry over 3 days. One hundred percent growth is defined as the highest parasitemia on the final day of the experiment. Data was fit to an exponential growth curve equation. Each data point is representative of the mean of 3 replicates ± S.E.M.

To visualize the PfGRP170-PfBiP interaction within the cellular context of the infected RBC, we utilized a Proximity Ligation Assay (PLA)^47–49^. The PLA positive signal indicates that two proteins are within 40nm of each other, suggesting a close interaction within the cell. This approach has been used successfully in *Plasmodium* to demonstrate interaction of exported proteins^50^. We performed this assay using anti-GFP and anti-BiP antibodies and observed a positive signal at all life cycle stages (Figure 5C). As a negative control we also probed with an antibody against the ER localized protease PMV and despite the co-localization of these two proteins in the ER, we did not see a positive PLA signal, suggesting distinct sub-organellar localizations (Figure 5D). Together, these results demonstrate that PfGRP170 and PfBiP interact during all stages of the asexual life cycle of *P. falciparum*.

The function of BiP is critical for ER biology and in other eukaryotes its function is regulated by GRP170^26, 27^. Additionally, loss of the PfGRP170 yeast homolog, Lhs1, activates a stress response mechanism known to upregulate BiP expression^51^. Therefore, we tested whether the PfGRP170-GFP-DDD mutants could be rescued by overexpression of PfBiP. We did this by episomally expressing PfBiP with a Ty1 tag and an ER retention signal (KDEL) in the PfGRP170-GFP-DDD mutants. Colocalization assays demonstrate that the PfBiP-Ty1 fusion protein is targeted to the ER and we observe that the protein is expressed at the expected size by western blot (Figure 5E and 5F). To determine if the overexpression of PfBiP could rescue parasite growth during TMP removal, the PfGRP170-GFP-DDD parasites expressing the PfBiP-Ty1 protein were incubated with and without TMP and the parasitemia was monitored using flow cytometry. We demonstrate that the overexpression of PfBiP in the PfGRP170-GFP-DDD parasites could not rescue parasite growth (Figure 5G).

### Loss of PfGRP170 function activates the PK4 stress response pathway

In addition to their function in the secretory pathway, molecular chaperones perform a vital role in the management of cellular stress. *Plasmodium* lack much of the ER machinery used to activate stress response pathways^16,52,53^. The only identified ER stress response pathway in *Plasmodium* is the PERK/PK4 pathway^10,16^. Signaling through this pathway has been shown to occur in the parasite following artemisinin treatment^10^. Under normal conditions, PK4 exists as a transmembrane monomeric protein in the ER. When the ER is stressed, PK4 oligomerizes and becomes active, phosphorylating the cytoplasmic translation initiation factor EIF2-α to halt translation and flux through the ER^10,16^. To determine whether this pathway was activated during conditional inhibition of PfGRP170, PfGRP170-GFP-DDD parasites were tightly synchronized to the ring stage and grown without TMP for 24 hours, after which parasite lysate was collected and the phosphorylation state of EIF2-α determined by western blot. We observed that PfGRP170 auto-inhibition resulted in the phosphorylation of EIF2-α, indicating that this pathway was activated (Figure 6A).

**Figure 6:**
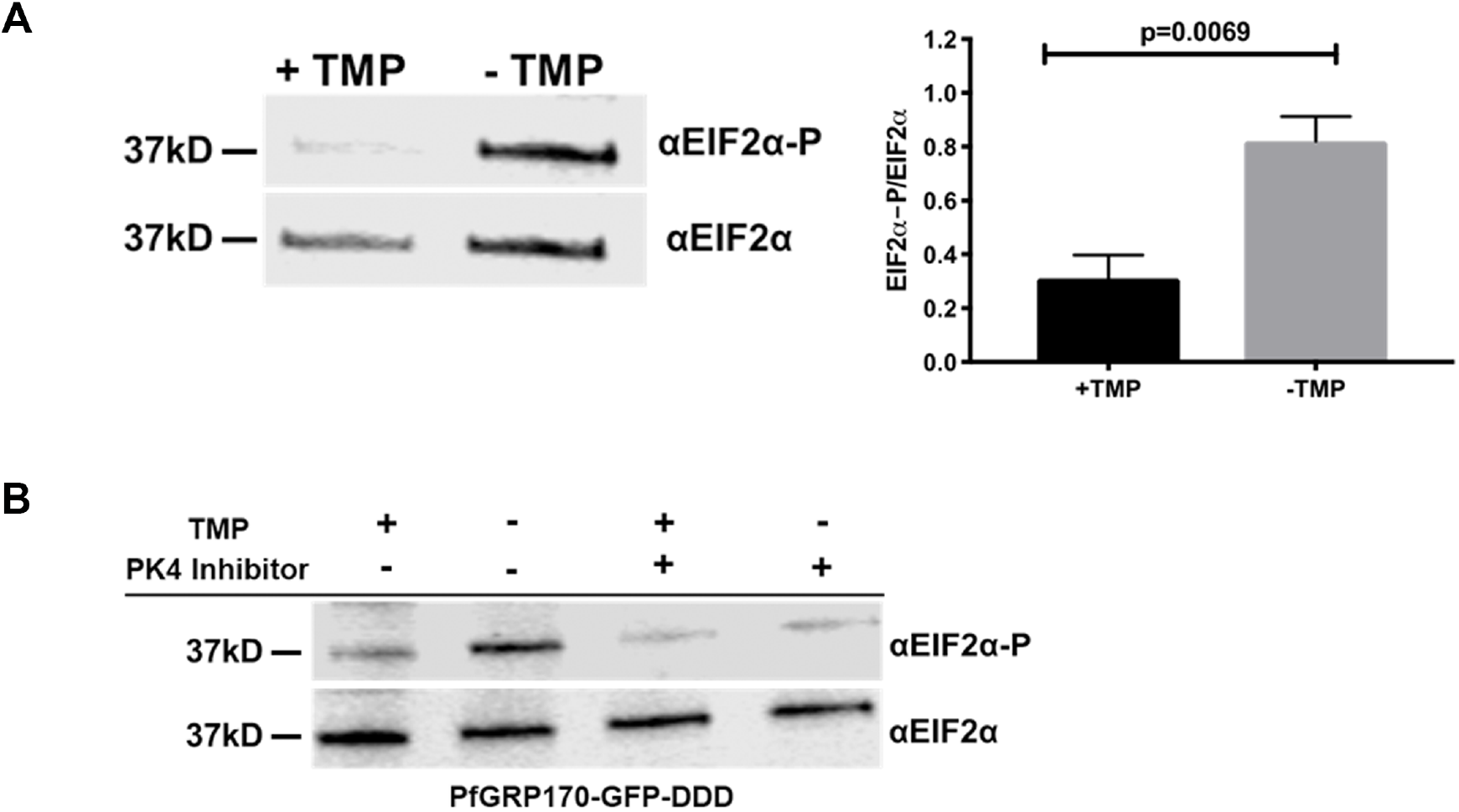
Loss of PfGRP170 function activates the PK4 stress pathway. **(A). (Left)**Synchronized ring stage PfGRP170-GFP-DDD parasites were incubated with and without TMP for 24 hours. Protein was isolated from these samples and analyzed via western blot, probing for anti-eIF2α and anti-Phospho-eIF2α. **(Right)** The ratio of phosphorylated EIF2α over total EIF2α for PfGRP170-GFP-DDD parasites incubated with and without TMP is shown. Western blot band intensities were calculated using ImageJ software (NIH) and the significance was calculated using an unpaired t test. Data are representative of 4 biological replicates ± S.E.M. **(B).** Synchronized ring stage PfGRP170-GFP-DDD parasites were incubated with and without TMP and in the presence and absence of 2μM PK4 inhibitor GSK2606414 for 24 hours. Protein was isolated from these samples and analyzed via western blot by probing for anti-eIF2α and anti-Phospho-eIF2α.

Since conditional inhibition of PfGRP170 resulted in EIF2-α phosphorylation, which has been shown to be required for resistance to artemisinin resistance, we tested if PfGRP170 plays a role in drug resistance. For this purpose, we utilized PfGRP170-BirA parasites, which have an extra copy of PfGRP170. Using the ring-stage survival assay, we compared the growth of the parental parasite line (3D7) with that of PfGRP170-BirA parasites after brief exposure to artemisinin. Our data show that overexpression of PfGRP170 did not result in artemisinin resistance (Supplemental Figure’s 8A-B).

Several *Plasmodium* kinases have been shown to phosphorylate EIF2-α in late developmental stages or in response to other cellular stress or artemisinin treatment^16,54–56^. We were therefore interested in identifying the specific kinase that is responsible for the phosphorylation of EIF2-α during conditional inhibition of PfGRP170. The ER kinase, PK4, has been shown to be activated by ER stress in *Plasmodium^53^*. Therefore, we incubated synchronized PfGRP170-GFP-DDD parasites without TMP for 24 hours, in the presence or absence of a specific PK4 inhibitor GSK2606414^10^. Parasite lysates were used to determine the phosphorylation state of EIF2-α. We observed that in the presence of the PK4 inhibitor, EIF2-α phosphorylation was blocked, demonstrating that conditional inhibition of PfGRP170 specifically results in PK4 activation, which leads to phosphorylation of EF2-α (Figure 6B). As a control, we used the parental strain, PM1, and incubated these parasites with and without TMP or the PK4 inhibitor (Supplemental Figure 9). This experiment showed no changes in levels of EIF2-α regardless of the presence of TMP or the PK4 inhibitor.

## DISCUSSION

We present in this work the first characterization of PfGRP170 in the asexual life cycle of *P. falciparum*. We have generated conditional mutants that allow us to probe the role of this protein using the DDD conditional auto-inhibition system^32,33,35–38^. Additionally, taking advantage of the GFP fused to PfGRP170, we were able to isolate an exceptionally rare clonal population using flow cytometry. This technique achieved what a traditional limiting dilution method could not. Moreover, this type of flow sorting can be implemented not only for rare events but also to significantly cut down the time from transfection to a clonal cell population.

We demonstrate here that PfGRP170 is an ER resident protein that is essential for asexual growth in *P. falciparum*. Loss of PfGRP170 function leads to a growth arrest of parasites late in development and their subsequent death. In yeast and mammals, GRP170 functions in a complex with the ER chaperone BiP, serving as the nucleotide exchange factor to regulate BiP activity^26,27^. Unlike *Plasmodium falciparum*, Yeast null for GRP170 are viable due to the upregulation of Sil1, another nucleotide exchange factor, that usually plays a role in the IRE1 stress response pathway^51^. The *Plasmodium* genome does not encode Sil1 and IRE1, which aligns with the observed essentiality of PfGRP170 during the blood stages. Additionally, research in mammalian systems suggests that GRP170 also has BiP-independent functions, such as binding unfolded substrates^28^. Our data show, via immunoprecipitation, mass spectroscopy, and proximity ligation assays, that PfGRP170 interacts with BiP in *P. falciparum* suggesting that it regulates BiP function. Further, overexpression of PfBiP was unable to rescue loss of PfGRP170 function and the conditional inhibition of PfGRP170 does not reduce its interaction with PfBiP. These data suggest that a PfBiP independent function of PfGRP170 is essential for parasite survival.

Previously it was shown that apicoplast transit peptides are predicted to bind the ER chaperone BiP, and when these predicted binding sites were mutated, targeting to the apicoplast was disrupted^57^. Moreover, an Hsp70 inhibitor with an antimalarial activity was shown to inhibit apicoplast targeting^58,59^. These data, combined with the predicted transit peptide of PfGRP170, led us to investigate the role of this chaperone in apicoplast trafficking. Interestingly, when the putative transit peptide was tagged with a GFP reporter and without an ER retention signal, the fusion protein was retained in the ER. It was previously reported that proteins with a signal peptide and no ER retention signal are secreted to the parasitophorous vacuole^60–63^. However, it was also shown that some proteins with a signal peptide and GFP (lacking an ER retention or trafficking signals) remain in the parasite ER^61^. Regardless, this reporter was not sent to the apicoplast indicating that it is not a functional apicoplast transit peptide. Previous work suggest that appending the first 137 amino acids of PfGRP170 to a GFP reporter (without a retention signal) resulted in this chimeric protein localizing partially to the apicoplast and to the parasitophorous vacuole^64^. Our chimeric protein includes the first 150 amino acids of PfGRP170 which may account for some of the differences in the two studies. In addition, PfGRP170 auto-inhibition did not lead to any defects in trafficking to the apicoplast, nor could it be rescued with the essential apicoplast metabolite IPP. Further, we did not identify any apicoplast localized proteins as potential interactors of PfGRP170 These data suggest that the primary function of PfGRP170 does not function in the apicoplast trafficking pathway.

Protein trafficking to the host RBC originates in the parasite ER and is essential for parasite viability, and therefore could potentially account for the observed death phenotype during conditional inhibition of PfGRP170^11,12^. PfGRP170 was shown to associate with exported proteins in another study that identified proteins that bind to the antigenic variant surface protein, PfEMP1^65^. However, our data show that there is no significant difference in the trafficking of some exported proteins upon conditional inhibition of PfGRP170, suggesting that protein export is not blocked.

ER chaperones are known in other eukaryotes to be vital to managing cellular stress^17,21^. However, several ER localized stress response pathways present in other eukaryotes are absent in *P. falciparum* and few molecular players in the parasite ER stress response pathway are known. Our data demonstrate that PfGRP170 is important for coping with a specific form of cellular stress, namely heat shock. This finding highlights a potential critical role for PfGRP170 *in vivo*, as high febrile episodes are one of the main symptoms of clinical malaria and are considered a defense mechanism against parasites. GRP170 in mammalian systems has been shown to bind to the transmembrane proteins in the ER that are involved in the unfolded protein response (UPR), suggesting it may regulate these pathways^66,67^. The *Plasmodium* genome does not encode many of the UPR orthologues, but a single ER stress pathway (PK4 signaling) has been previously described and was shown to be activated following artemisinin treatment^10,16^. Here, we demonstrate that loss of PfGRP170 function results in the activation of PK4 stress pathway, providing the first link between an endogenous ER resident protein and the activation of the PK4 pathway in *P. falciparum*. Further, our data suggest that even though the PK4 stress response pathway is activated upon removal of TMP, this pathway is ultimately unable to prevent parasite death. This is most likely because one or more essential proteins that depend of PfGRP170 for their correct folding and function.

Yeast null for the GRP170 homolog, Lhs1, activate the IRE1 UPR signaling pathway^51^. Activation of the IRE1 pathway in eukaryotes, which is not present in *Plasmodium*, typically leads to the upregulation of ER chaperones such as BiP^51–53,68^. These data suggest that the only essential function of Lhs1 is to serve as a nucleotide exchange factor for BiP^51^. Therefore, we tested whether the death phenotype seen in the conditional PfGRP170 mutants could be rescued by overexpressing PfBiP. Our experiments revealed that overexpression of PfBiP does not improve viability of the PfGRP170-GFP-DDD parasites following TMP removal. This data implies, that unlike its homologs in other eukaryotes, the essential function of PfGRP170 may not be entirely linked to its role in regulating BiP.

We utilized two separate IP/mass spectroscopy approaches to generate a list of 11 high-confidence interacting partners of PfGRP170. Seven of the proteins (including PfBiP) have a peak expression pattern around the time PfGRP170-GFP-DDD parasites begin to die. SERA5 and SERA6 have been shown to be required for egress from the RBC, which would be after the PfGRP170-GFP-DDD parasites die^69–71^. RON3 has been shown to been suggested to be a protein important for RBC invasion, which implies this protein interaction is also not why PfGRP170-GFP-DDD parasites are dying^72^. CLAG9, another identified protein, has been proposed to play a role in cytoadherence to CD36 and remodeling the host RBC after invasion by a merozoite^73,74^. PDI-11 was predicted to be non-essential in a *piggyBac* mutagenesis conducted in *Plasmodium^75^*. The remaining protein was parasite-infected erythrocyte surface protein 1 (PIESP1). Overexpression data suggest that PIESP1 is exported to the host RBC^76^. However, this protein has a putative ER retention signal (TDEL). These last four amino acids were left off of the GFP fusion protein that was expressed in the parasite as the authors predicted this protein was a transmembrane protein and thus leaving off these amino acids would have no effect on protein localization^76^. Further studies will be needed to determine the precise subcellular localization of PIESP1 and determine its role in parasite biology. These data show that PfGRP170 is essential for the asexual lifecycle of *P. falciparum* and that the biological role of PfGRP170 is quite divergent from other eukaryotes. Further, given the divergence between mammalian and parasite GRP170s, PfGRP170 could be a viable antimalarial drug target.

---

**Supplemental Figure 1:**
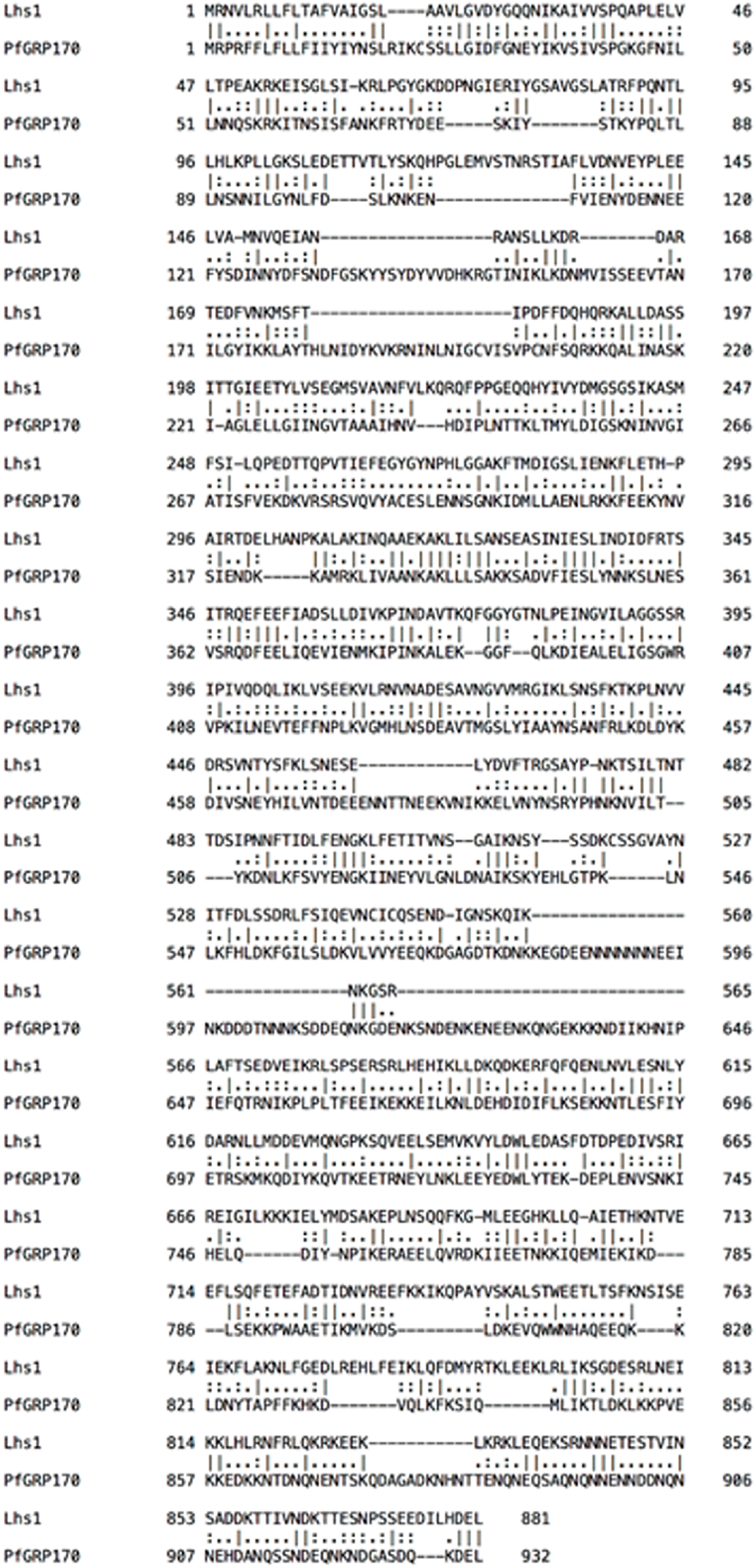
Sequence Alignment of Lhs1 and PfGRP170. Sequence alignment of *S. cerevisiae* GRP170 (Lhs1) and PfGRP170. The alignment was performed using EMBOSS Needle which creates a global alignment of two sequences using the Needleman-Wunsch algorithm. The software used to do this is provided by the European Bioinformatics Institute, which is a part of the European Molecular Biology Laboratory (EMBL). Identical residues are indicated by a “I”, strongly similar residues are indicated by a “:”, and weakly similar residues are indicated by a “.”.

**Supplemental Figure 2:**
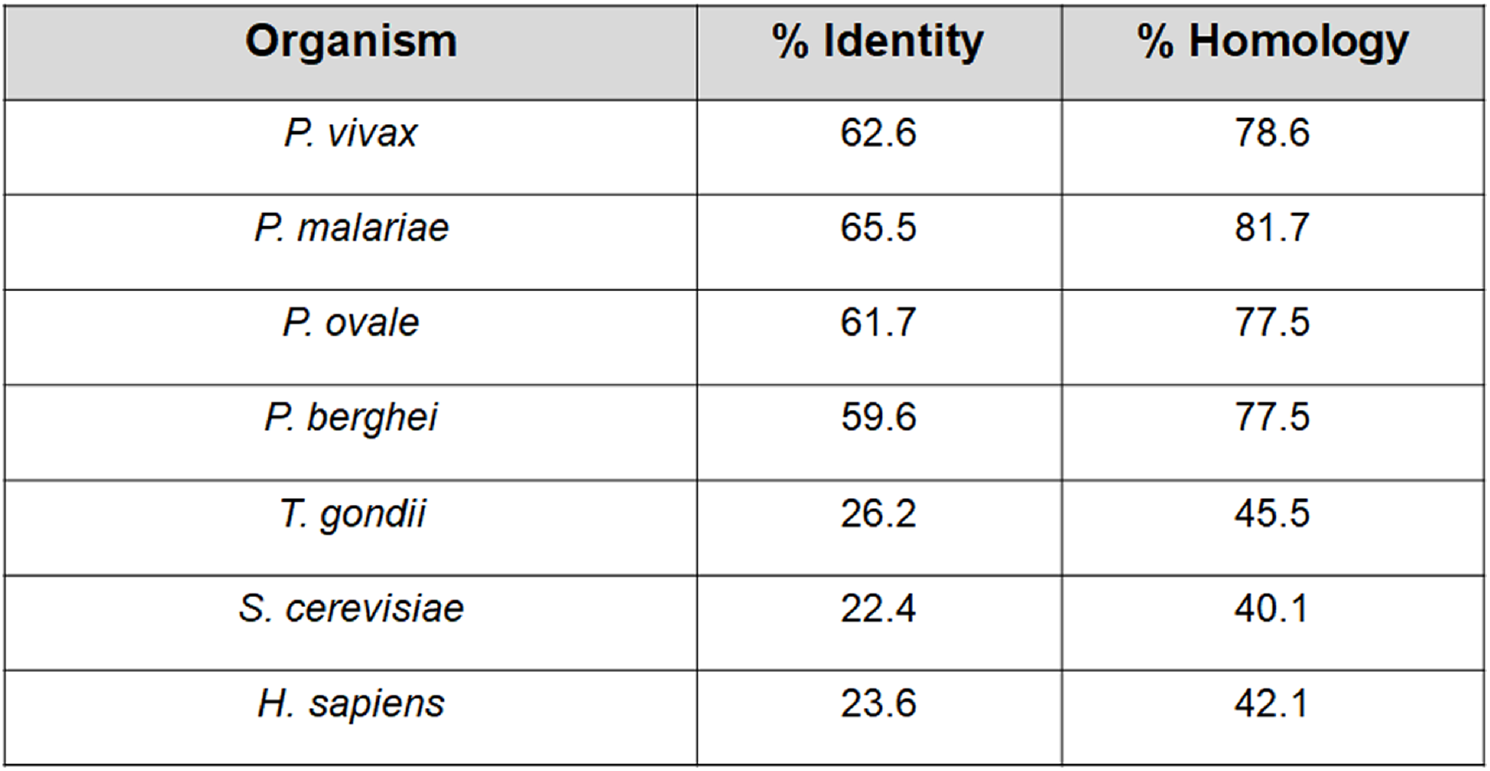
Sequence homology of PfGRP170. Sequence identify and homology of *P. falciparum* GRP170 compared to GRP170 homologs from other *Plasmodium* Species (*P. vivax* (PVX_083105), *P. malariae* (PmUG01_12020700), *P. ovale* (PocGH01_12018900), and *P. berghei* (PBANKA_1357200)), *T. gondii* GRP170 (TGGT1_226830), yeast GRP170 (*S. cerevisiae*), and human GRP170 (*H. sapiens*). Alignments to determine sequence identify and homology were performed using EMBOSS Needle which creates a global alignment of two sequences using the Needleman-Wunsch algorithm. The software to do this is provided by the European Bioinformatics Institute, which is a part of the European Molecular Biology Laboratory (EMBL).

**Supplemental Figure 3:**
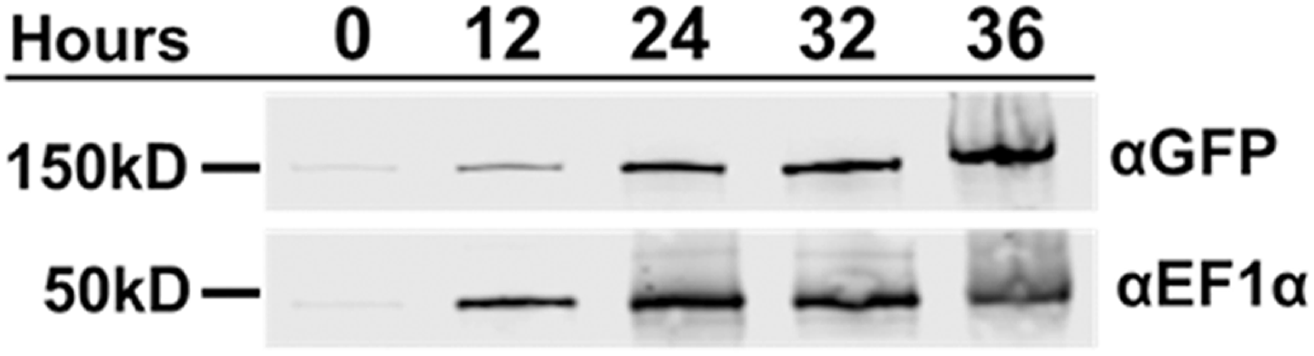
PfGRP170 is Expressed Throughout the Asexual Life Cycle. TMP was removed from tightly synchronized ring stage PfGRP170-GFP-DDD parasites and protein was isolated throughout the asexual life cycle. Lysates were separated on a Western blot and probed with anti-GFP to visualize PfGRP170-GFP-DDD and anti-PfEF1α as a loading control.

**Supplemental Figure 4:**
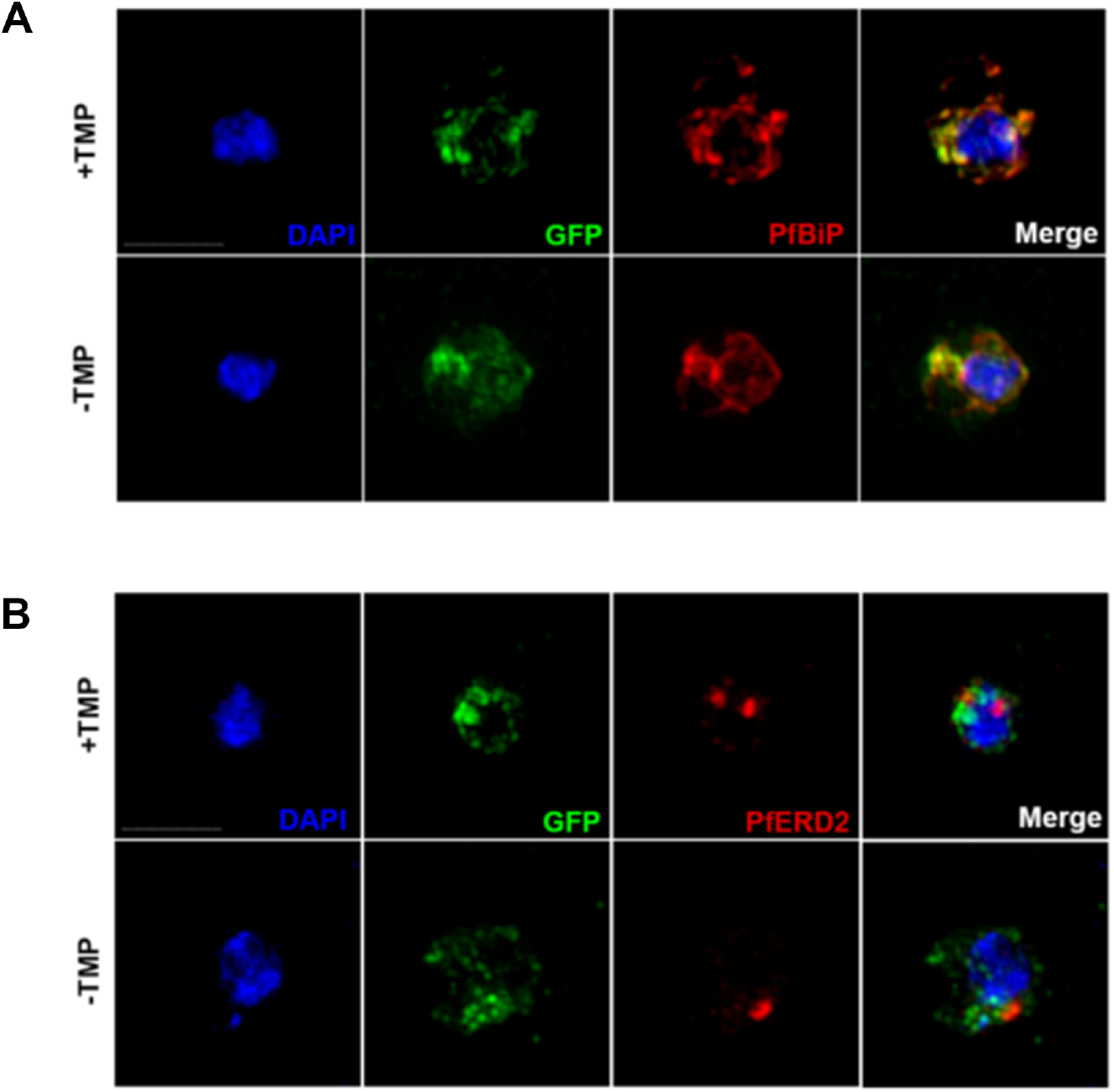
Conditional mutants of PfGRP170 localize to the ER. Synchronized PfGRP170-GFP-DDD ring stage parasites were incubated with and without TMP for 24 hours. Parasites were then fixed with paraformaldehyde and stained with either DAPI, anti-GFP, and anti-BiP (ER) **(A)** or DAPI, anti-GFP, and anti-ERD2 (Golgi) **(B)**. Images were taken as a Z-stack using super resolution microscopy and SIM processing was performed on the Z-stacks. Images are displayed as a maximum intensity projection. The scale bar is 2μm.

**Supplemental Figure 5:**
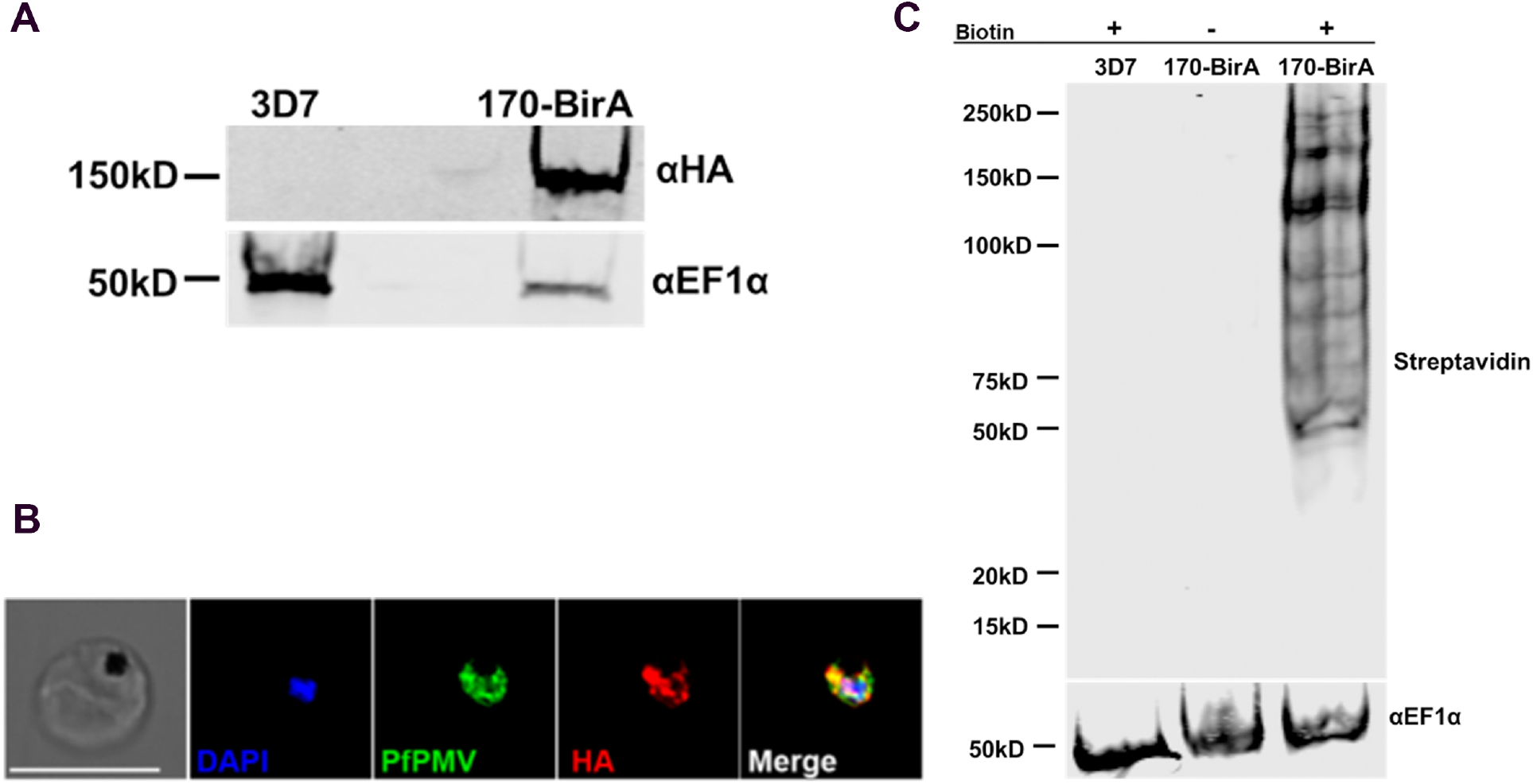
PfGRP170-BirA localizes to the parasite ER and biotinylates proteins. **(A).** Western blot of 3D7 (parental) and PfGRP170-BirA expressing parasites probed with anti-HA and anti-EF1α. **(B).** Paraformaldehyde fixed PfGRP170-BirA parasites stained with anti-HA (PfGRP170-BirA), anti-PfPMV (ER), and DAPI. The images were taken with Delta Vision II, deconvolved and are displayed as a maximum intensity projection. The scale bar is 5μm. **(C).** A western blot analysis of 3D7 (parental) and PfGRP170-BirA parasites following a 24-hour incubation with biotin is shown. A fluorophore-labeled streptavidin secondary antibody was used to visualize biotinylated proteins. A control with PfGRP170-BirA parasites incubated without biotin is also shown. Anti-EF1α is used as a loading control.

**Supplemental Figure 6:**
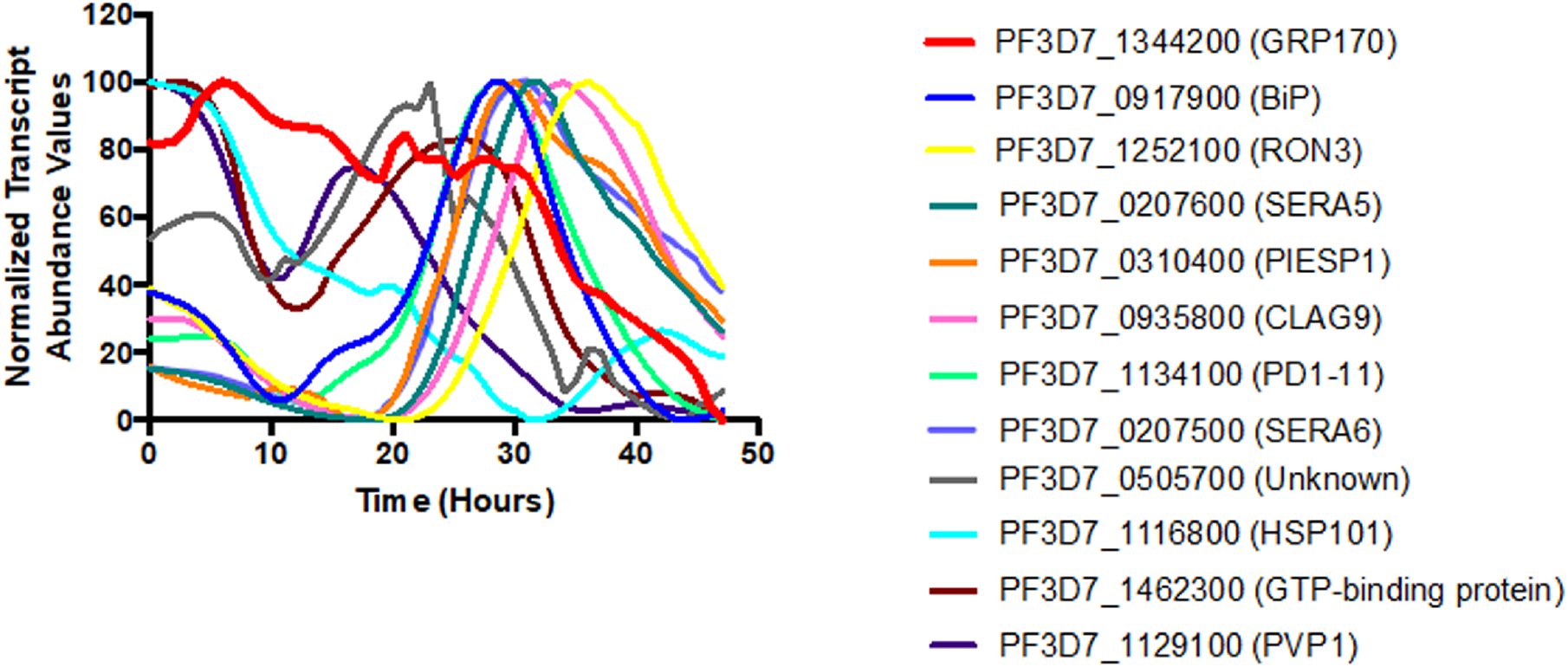
Relative transcript abundance of proteins identified in both the anti-GFP co-immunoprecipitation and BioID mass spectroscopy approaches. The relative transcript abundance of the 11 PfGRP170 interacting proteins identified in Figure 4. The data are plotted using previously published genome-wide real-time transcription data^46^.

**Supplemental Figure 7:**
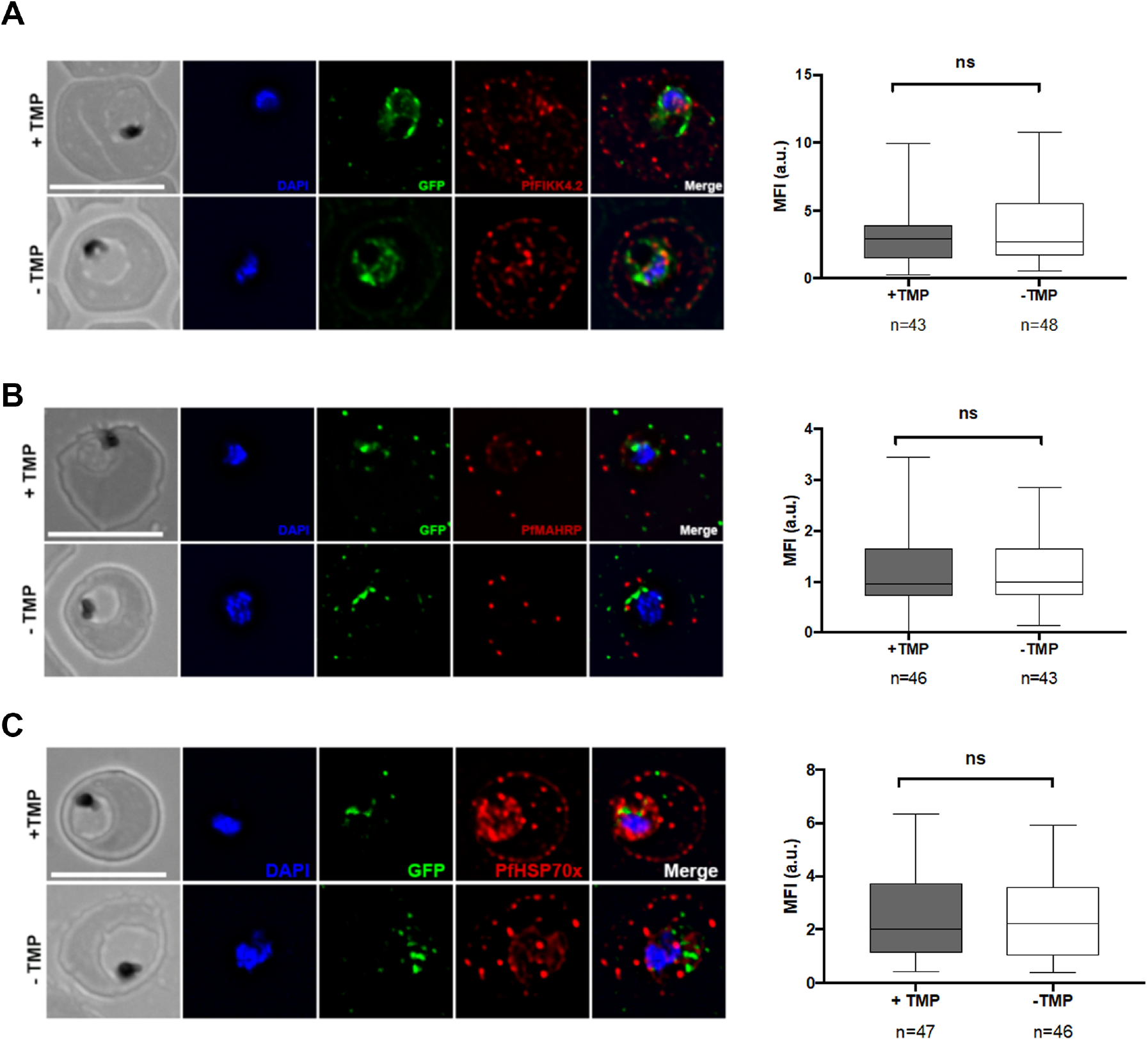
PfGRP170 is not Required for Trafficking to the Host RBC. Tightly synchronized ring stage PfGRP170-GFP-DDD parasites were incubated with and without TMP for 24 hours. Following this incubation, parasites were fixed with acetone and stained with DAPI, anti-GFP (PfGRP170) and either anti-PfFIKK4.2 **(A),** anti-PfMAHRP1C **(B),** or anti-PfHSP70X **(C).** The images were taken with Delta Vision II, deconvolved, and are displayed as a maximum intensity projection. The scale bar is 5μM. Mean Fluorescent Intensity (M.F.I) was calculated for the exported fraction (PfFIKK4.2, PfMAHRP1C, and PfHSP70x) from individual cells. Data are from two independent experiments and is displayed as box-and-whiskers plots (whiskers represent the maximum and minimum M.F.I). The significance was calculated using an unpaired t test (NS= not significant).

**Supplemental Figure 8:**
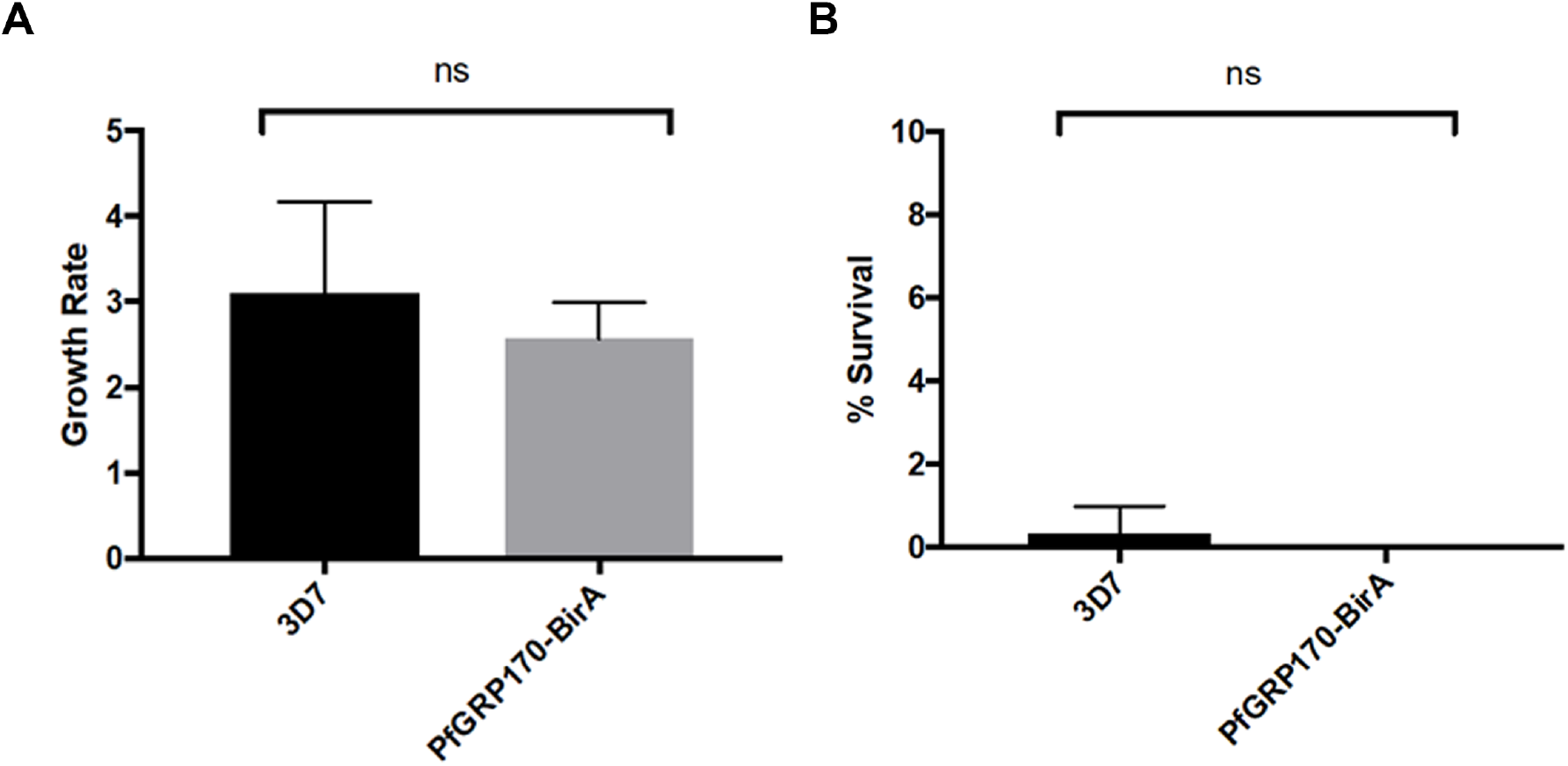
Overexpression of PfGRP170 does not Confer Artemisinin Resistance. Tightly synchronized ring stage 3D7 and PfGRP170-BirA parasites were incubated with either 1% DMSO (Control) or Dihydroartemsinin (DHA) for 6 hours. After 6 hours the drug is removed by washing the culture with complete RPMI. Parasitemia was calculated using Giemsa stained thin blood smears at 0 hours (to calculate starting parasitemia) and 72 hours after either DMSO or DHA exposure. Four independent replicates of the experiment were completed for 3D7 and three for PfGRP170-BirA. The growth rate of the 3D7 and PfGRP170-BirA parasites, incubated only with DMSO, was calculated after 72 hours **(A).** The percent survival of parasites was calculated for 3D7 and PfGRP170-BirA after DHA exposure was calculated after 72 hours **(B).**

**Supplemental Figure 9:**
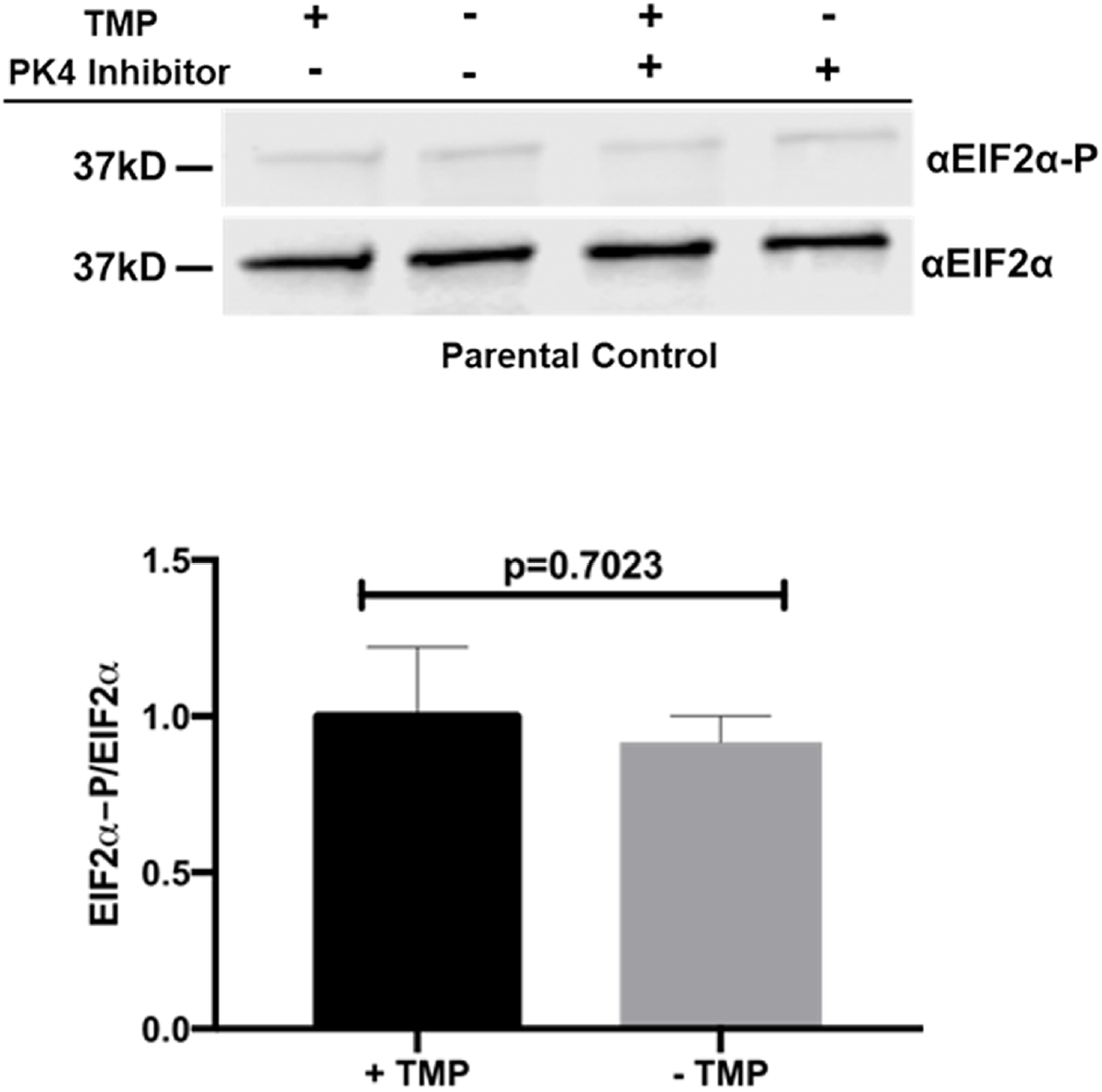
EIF2-α levels do not change in PM1 parasites in the presence or absence of TMP or a PK4 inhibitor. **(Top)** Synchronized ring stage PM1 parasites were incubated with and without TMP and in the presence and absence of 2μM PK4 inhibitor GSK2606414 for 24 hours. Protein was isolated from these samples and analyzed via western blot by probing for anti-eIF2α and anti-Phospho-eIF2α. **(Bottom)** The ratio of phosphorylated EIF2α over total EIF2α in PM1 parasites incubated with and without TMP is shown. Western blot band intensities were calculated using ImageJ software (NIH) and the significance was calculated using an unpaired t test. Data are representative of 2 biological replicates ± S.E.M.

**Supplemental Table 1: Raw Mass Spectroscopy data**

Raw mass spectroscopy data from two anti-GFP IP (using 1B2 and 1B11), two parental anti-GFP IP’s (using PM1), two Streptavidin IP’s the PfGRP170-BirA cell lines following a 24 incubation with biotin, and one streptavidin IP’s of 3D7 cell lines following a 24 hour incubation with biotin. Both approaches used asynchronous cells. The excel file includes the following: A total list of proteins from PlasmoDB containing a signal peptide and/or transmembrane domain used to sort the mass spectroscopy data (Tab 1), raw mass spectroscopy data from PM1 anti-GFP IP’s 1 and 2 (Tabs 2 and 3), list of proteins from the PM1 anti-GFP IP’s 1 and 2 that contained a signal peptide and/or transmembrane domain (Tabs 4 and 5), raw mass spectroscopy data from the 1B2 anti-GFP IP (Tab 6) and the list of proteins from the 1B2 anti-GFP IP that contained a signal peptide and/or transmembrane domain (Tab 7), raw mass spectroscopy data from the 1B11 anti-GFP IP (Tab 8) and the list of proteins from the 1B11 anti-GFP IP that contained a signal peptide and/or transmembrane domain (Tab 9), raw mass spectroscopy data from a streptavidin IP on 3D7 parasites incubated with biotin (Tab 10), the list of proteins from the 3D7 streptavidin IP that contained a signal peptide and/or transmembrane domain (Tab 11), raw mass spectroscopy data from two independent PfGRP170-BirA streptavidin IP’s (Tabs 12 and 13) and the list of proteins from the PfGRP170-BirA streptavidin IP’s that contained a signal peptide and/or transmembrane domain (Tabs 14 and 15).

## ACKNOWLEDGMENTS

We thank Dan Goldberg for anti-EF1α and anti-PMV antibodies; Boris Striepen for anti-Cpn60 antibody; Hans-Peter Beck for anti-MAHRP antibody; Jude Przyborski for anti-PfHSP70x antibody; David Cavanagh and EMRR for anti-FIKK4.2 antibody; Drew Etheridge, Min Zhang, and Bill Sullivan for technical suggestions; Muthugapatti Kandasamy at the University of Georgia Biomedical Microscopy Core, Julie Nelson at the CTEGD Cytometry Shared Resource Lab for technical assistance. We acknowledge assistance of the Emory University Integrated Proteomics Core for mass spectrometry. This work was supported by ARCS Foundation awards to H.M.K. and D.W.C., UGA Startup funds to V.M., CDC-UGA Seed Award to V.M. and N.W.L., and the US National Institutes of Health (R00AI099156 and R01AI130139) to V.M. and (T32AI060546) to H.M.K. and to M.A.F.

## METHODS

### Primers and Plasmid construction

All primer sequences used in this study can be found in Supplemental Table 2.

Generation of pGDB-SDEL plasmid was done using the QuikChange II Site-Directed Mutagenesis Kit (Agilent Technologies) on the pGDB plasmid with primers P1 and P2 per the manufacturer’s protocol^36^.

Genomic DNA was isolated using the QIAamp DNA blood kit (Qiagen). gDNA used in this study was isolated from either 3D7 or Plasmepsin I knockout parasites (PM1KO)^36^. The pPfGRP170-GFP-DDD plasmid used to generate the PfGRP170-GFP-DDD mutants was made by amplifying via PCR an approximately 1-kb region homologous to the 3’end of the PfGRP170 gene (stop codon not included) using primers P3 and P4. The amplified product was inserted into pGDB-SDEL plasmid using restriction sites Xho1 and AvrII (New England Biolabs) and transformed into bacteria. The construct was sequenced prior to transfection.

The pGRP170-HA-BirA-KDEL plasmid was prepared by amplifying PfGRP170 (without the stop codon) from 3D7 gDNA using primers P5 and P6 and 3xHA-BirA from the pTYEOE-3XHA-BirA plasmid (From D. Goldberg) using primers P7 and P8. Both PCR products generated included homologous regions used for Sequence and Ligation Independent Cloning (SLIC)^77^. The primers to amplify the 3xHA-BirA included the sequence of an ER retention signal (KDEL). These PCR products were fused together using PCR sewing as described previously and subsequently PCR amplified using primers P5 and P8^35^. The resulting product was then inserted into pCEN-DHFR^78^ that was digested with Nhe1 and BglII (New England Biolabs) using SLIC and transformed into bacteria as described previously^32,35^.

The pPfGRP170TP-GFP plasmid was prepared by amplifying the first 450 bp (includes the signal peptide and putative transit peptide sequence) of PfGRP170 from PM1 gDNA using primers P5 and P9. The GFP sequence used was amplified from pGDB using primers P10 and P11. The PfGRP170 transit peptide PCR was digested with Nhe1 and AatII (New England Biolabs) and the GFP PCR was digested with AatII and BglII (New England Biolabs). The two fragments were then ligated together (via the AatII digest site) using a T4 ligase (kit from New England Biolabs) and subsequently PCR amplified using primers P5 and P11. The resulting product was then digested with Nhe1 and BglII and inserted into pCEN-DHFR^78^ that was digested with Nhe1 and BglII (New England Biolabs) using a T4 ligase and transformed into bacteria as described previously^32,35^.

The pPfBiP-Ty1overexpression plasmid was prepared first by generating cDNA using the SuperScript III reverse transcriptase kit (Invitrogen) using primer P14. PfBiP was then amplified from the cDNA using primers P14 and P15. The resultant PCR product included PfBiP, a single Ty1 tag, and an ER retention signal (KDEL). The pCEN vector was modified to contain the DHOD resistance gene instead of the DHFR for parasite selection^78^. The PfBiP-Ty1-KDEL-KDEL PCR was cloned into the pCEN-DHOD vector cut with Nhe1 and BglII (New England Biolabs) using the IN-Fusion HD EcoDry Cloning Kit (Clontech).

### Cell Culture, transfections, and isolation of clonal cell lines

Parasites were grown in RPMI 1640 media supplemented with Albumax 1 (Gibco) and transfected as described previously^31–33,35,36^.

To generate PfGRP170-GFP-DDD mutants, PM1KO parasites were transfected with the pPfGRP170-GFP-DDD plasmid in duplicate. PM1KO parasites contain the human dihydrofolate reductase (hDHFR) expression cassette which gives the parasites resistance to Trimethoprim (TMP)^36^. Drug selection and cycling were performed as described previously using 10μM TMP (Sigma) and 2.5μg/ml Blasticidin (Sigma)^32,33,36^. Following drug cycling, GFP positive cells were enriched using an S3 Cell Sorter (BioRad). Individual GFP positive cells from a single transfection were cloned into 96 well plates using a MoFlo XDP flow cytometer. After the EC_50_ of TMP was determined for clones 1B2 and 1B11, parasites were shifted into media containing 2.5μg/ml BSD and 20μM TMP to facilitate optimal growth.

The PfGRP170-BirA and PfGRP170TP-GFP parasites were generated by transfecting 3D7 parasites with plasmids pGRP170-HA-BirA-KDEL or pPfGRP170TP-GFP, respectively. Parasites expressing these episomal constructs were selected using 2.5nM WR99210.

To generate the PfGRP170-GFP-DDD parasites episomally expressing PfBiP-Ty1-KDEL-KDEL, PfGRP170-GFP-DDD clones 1B2 and 1B11 were each transfected with pPfBiP-Ty1-KDEL. This plasmid expresses PfBiP-Ty1-KDEL using the *pbef1α* bi-directional promoter. Parasites expressing this episomal construct were selected using 250nM of DSM1^79^.

### Integration tests for PfGRP170-GFP-DDD mutants

Genomic DNA was isolated from parasites using the QIAamp DNA blood kit (Qiagen). Control primers to amplify the genome were P4 and P12 and primers used to amplify integrated DNA were P12 and P13.

Southern blot analysis was performed on DNA isolated from PfGRP170-GFP-DDD parasites (1B2 and 1B11) as described previously^32,35^. The assay was also performed on PM1KO parental DNA and the pGRP170-DDD plasmid as a control. DNA was isolated from parasites using the QIAamp DNA blood kit (Qiagen). 10μg of precipitated PM1KO DNA, 1B2, and 1B11 DNA and 10ng of pGRP170-DDD plasmid was digested overnight with Mfe1 (New England Biolabs). The biotinylated probe used was generated by PCR using biotinylated-16-UTP (Sigma) and primers P3 and P4. The biotinylated probe on the southern blot was detected using IRDye 800CW streptavidin-conjugated dye (LICOR Biosciences) and imaged using the Odyssey infrared imaging system (LICOR Biosciences).

### Growth assays using flow cytometry

TMP was removed from asynchronous PfGRP170-GFP-DDD cultures for growth assays by washing the culture in equal volume of complete RPMI three times. The culture was then resuspended in complete RMPI media containing either 2.5μg/ml Blasticidin (Sigma) for conditional inhibition (Sigma) or 2.5μg/ml Blasticidin (Sigma) and 20μM TMP (Sigma) for the control. Parasitemia was monitored using a flow cytometer, either a CyAn ADP (Beckman Coulter) or CytoFLEX (Beckman Coulter) instrument, using either 1.5μg/ml acridine orange (Molecular Probes) as described previously^35^ or similarly using 8μM Hoechst in filtered 1X phosphate-buffered saline (PBS). Flow cytometry data were analyzed using FlowJo software (Treestar Inc.). If the parasitemia was too high, parasites were subcultured during the experiment and the relative parasitemia was then calculated by multiplying the calculated parasitemia by the dilution factor. Parasitemia was normalized by using the highest parasitemia as one hundred percent. Using Prism software (GraphPad Software Inc), the parasitemia data were fit to an exponential growth curve equation.

To determine the EC_50_ of TMP for PfGRP170-GFP-DDD cell lines, parasites were washed as described above and seeded into a 96 well plate with 2.5μg/ml Blasticidin and varying TMP concentrations. Parasitemia was measured after 48 hours using flow cytometry as described above. The parasitemia data were fit to a dose-response equation using Prism. For the IPP rescue experiment, asynchronous PfGRP170-GFP-DDD parasites were washed as described above and resuspended in media either with 2.5μg/ml Blasticidin or 2.5μg/ml Blasticidin and 20μM TMP with or without 200μM Isopentenyl pyrophosphate (Isoprenoids LC). Parasitemia were monitored using flow cytometry as described above and the data were fit to an exponential growth curve equation using Prism.

For the heat shock experiment, asynchronous PfGRP170-GFP-DDD parasites were washed as described above and resuspended in media either with 2.5μg/ml Blasticidin or 2.5μg/ml Blasticidin and 20μM TMP. Parasites were then incubated at either 37°C or 40°C for 6 hours. After 6 hours, 20μM TMP was added to cultures that were incubated without it and all parasites were shifted back to 37°C. Parasitemia was monitored using flow cytometry as described above and the data were fit to an exponential growth curve equation (GraphPad Software Inc).

For growth assays done with PfGRP170-GFP-DDD-GFP cell lines overexpressing PfBiP, asynchronous parasites were washed as described above and resuspended in media either with 2.5μg/ml Blasticidin and 250nM of DSM1 or 2.5μg/ml Blasticidin, 250nM of DSM1, and 20μM TMP. Parasitemia were monitored using flow cytometry as described above and the data were fit to an exponential growth curve equation using Prism.

### Synchronized growth assay

PfGRP170-GFP-DDD Parasites were synchronized as described previously by sorbitol (VWR), followed by percoll (Genesee Scientific) the next day and then sorbitol four hours later to obtain 0-4 hour rings^32,36^. Parasites were washed as described above to remove TMP from the media and incubated in media either with 2.5μg/ml Blasticidin or 2.5μg/ml Blasticidin and 20μM TMP. Thin blood smears using the Hema 3 Staining Kit (PROTOCOL/Fisher) were prepared every few hours to monitor parasite growth and morphology. Slides were imaged using a Nikon Eclipse E400 microscope with a Nikon DS-L1-5M imaging camera.

### Western blot

Western blotting was performed as described previously^32^. Parasite pellets were isolated using cold 0.04% Saponin (Sigma) in 1X PBS for 10 minutes as described previously^32,36^. Antibodies used for this study were: mouse anti-GFP JL-8 (Clontech, 1:3000), rabbit anti-PfEF1α (from D. Goldberg, 1:2,000), mouse anti-plasmepsin V (from D. Goldberg, 1:400), rabbit anti-PfBiP MRA-1246 (BEI resources, 1:500), rabbit anti-GFP A-6455 (Invitrogen, 1:2,000), mouse anti-eIF2α L57A5 (Cell Signaling, 1:1,000), rabbit anti-Phospho-eIF2α 119A11 (Cell Signaling, 1:1,000), rat anti-HA (Roche 3F10, 1:3000), mouse anti-Ty1 (Sigma Clone BB2, 1:1000), and mouse anti-Ub P4D1 (Santa Cruz Biotechnology, 1:1,000). Secondary antibodies used were IRDye 680CW goat anti-rabbit IgG and IRDye 800CW goat anti-mouse IgG (LICOR Biosciences, 1:20,000). The western blots were imaged using the Odyssey infrared imaging system. Polyacrylamide gels used in this study were either prepared using 10% EZ-Run protein gel solution (Fisher) or precast gradient gels (4-20%, from Biorad). Any quantification performed on western blots was done using ImageJ software. The quantification data were analyzed using Prism (GraphPad Software, Inc.).

### PK4 Inhibitor Experiments

Synchronized ring stage PfGRP170-GFP-DDD parasites were incubated in media with either 2.5μg/ml Blasticidin or 2.5μg/ml Blasticidin and 20μM TMP in the presence or absence of a PK4 inhibitor GSK2606414 (Millipore Sigma) at 2μM for 24 hours. After 24 hours, the parasites were lysed for western blot analysis using 0.04% saponin in 1X PBS as described above. PM1 (parental control parasites) were incubated in media with either complete RPM1 (no drug) or media containing 20μM TMP in the presence or absence of PK4 inhibitor GSK2606414 (Millipore Sigma) at 2μM for 24 hours. After 24 hours, the parasites were lysed for western blot analysis using 0.04% saponin in 1X PBS as described above.

### Live Fluorescence Microscopy

To visualize PfGRP170-GFP-DDD live parasites, 100μL of parasite culture was pelleted. The supernatant was removed, and the parasites were resuspended in 100μL medium with 2.5μg/ml Blasticidin and 20μM TMP and 5μM Hoechst. The parasites were incubated at 37°C for 20 minutes. The parasites were then pelleted again and 90% of the medium was removed. Parasites were resuspended in the remaining medium and 8μL of this culture was placed on a glass slide and covered with a coverslip. The edges were sealed with nail polish and the cells were imaged using a DeltaVision II Microscope.

### Immunofluorescence trafficking assays and imaging processing

Immunofluorescence assays (IFA) were performed as described previously using a combination of 4% Paraformaldehyde and 0.015% glutaraldehyde for fixation and permeabilization using 0.1% Triton-X100^32,35^ or by smearing cells on a slide and fixing them with acetone. For apicoplast and red blood cell trafficking assays, cells were synchronized and TMP was removed as described above. Cells were then fixed as described above, 24 hours after the removal of TMP.

Primary antibodies used for IFAs in this study were: rabbit anti-GFP A-6455 (Invitrogen, 1:200), rat anti-PfBiP MRA-1247 (BEI resources, 1:125), rabbit anti-PfBiP MRA-1246 (BEI resources (1:100), mouse anti-plasmepsin V (From D. Goldberg, 1:1), mouse anti-GFP clones 7.1 and 13.1 (Roche 11814460001, 1:500), rabbit anti-Cpn60 (From. B. Striepen, 1:1,000), rabbit anti-PfERD2 (MR4, 1:2,000), rabbit anti-HA 9110 (Abcam, 1:200), rabbit anti-PfMAHRP1C (From. Hans-Peter Beck, 1:500), mouse anti-PfFIKK4.2 (From David Cavanagh/EMRR, 1:1,000), mouse anti-Ty1 (Sigma Clone BB2, 1:200), and rabbit anti-PfHSP70X (From Jude Przyborski, 1:500). Secondary antibodies used in this study are Alexa Fluor goat anti-rabbit 488, Alexa Fluor goat anti-rabbit 546, Alexa Fluor goat anti-mouse 488, Alexa Fluor goat anti-mouse 546, and Alexa Fluor goat anti-rat 546 (Life Technologies, 1:100). The mouse anti-PfFIKK4.2, rabbit anti-PfHSP70X, and anti-PfMAHRP1C require acetone fixation.

All fixed cells were mounted using ProLong Diamond with DAPI (Invitrogen) and imaged using the DeltaVision II microscope system or Zeiss ELYRA S1 (SR-SIM) Super Resolution Microscope using a 100X objective. Images taken using the DeltaVision II were collected as a Z-stack and were deconvolved using the DeltaVision II software (SoftWorx). The deconvolved Z-stacks were then displayed as a maximum intensity projection using SoftWorx. Images taken using the Super Resolution Microscope were taken as a Z-stack. The Z-stacks were analyzed using Zen Software (Zeiss, version from 2011) for SIM processing and obtaining the maximum intensity projection. Any adjustments made to the brightness and/or contrast of the images were made using either Softworx, Zen Software, or Adobe Photoshop and were done for display purposes only. Any quantification performed for microscopy images was done using ImageJ software as described previously^35^. The quantification data were analyzed using Prism (GraphPad Software, Inc.).

### Co-immunoprecipitation assays and Mass Spectroscopy

Parasites pellets were isolated from 48 mL of asynchronous culture at high parasitemia (10% or higher) using cold 0.04% saponin in 1X PBS as described above. Parasite pellets were lysed by resuspending the pellet in 150μL of Extraction Buffer (40mM Tris HCL pH 7.6, 150mM KCL, and 1mM EDTA) with 0.5% NP-40 (VWR) and 1X HALT protease inhibitor (Thermo). The resuspended parasites were then incubated on ice for 15 minutes and then sonicated three times (10% amplitude, 5 second pulses). In between each sonication, the lysate was placed on ice for 1 minute. The lysate was then centrifuged at 21,100g for 15 minutes at 4°C. The supernatant was collected in a fresh tube and placed on ice. The remaining pellet was subjected to a second lysis step using 150μL of Extraction buffer as above without NP-40. The lysate was sonicated and centrifuged as above (no 15-minute incubation on ice). The supernatant was collected and combined with the lysate from the first lysis step (INPUT sample). 20μL of the input sample was collected into a fresh tube and stored in the −80°C. The remaining input sample was combined with 2μL of rabbit anti-GFP monoclonal G10362 (Thermo) and incubated rocking for two hours at 4°C.

After the two-hour incubation, the lysate with antibody was added to 50μL of prepared protein G Dynabeads (Invitrogen). Dynabeads were prepared by washing 50μL of beads three times with 100μL of IgG binding buffer (20mM Tris HCL pH 7.6, 150mM KCL, 1mM EDTA, and 0.1% NP-40). The IgG binding buffer was removed from the beads each time using a magnetic rack (Life technologies). The beads, antibody, and lysate were incubated rocking for two hours at 4°C. After the two-hour incubation, the unbound fraction of protein was collected using the magnetic rack into a fresh tube and stored at −80°C until needed for western blot analysis. The beads were then washed two times in 300μL of IgG binding buffer with 1X HALT and one time in IgG binding buffer with 1X HALT without NP-40. Each wash was done for 10 minutes rocking at 4°C.

For Co-IP’s to show PfGRP170-GFP-DDD/BiP interaction 0-4 hour ring stage parasites were obtained and TMP was removed as described under the synchronized growth assay section. Parasites were lysed and an anti-GFP IP was performed as described above, approximately 24 hours after the removal of TMP. Protein was eluted off the beads for western blot using 1X Protein Loading Dye (LICOR) with 2.5% beta-Mercaptoethanol (Fisher) and boiled for 5 minutes. This was followed by a centrifugation at 16,200 g for 5 minutes. The eluted proteins are collected by placing the tube on a magnetic rack. The isolated proteins on magnetic beads were digested with trypsin and analyzed at the Emory University Integrated Proteomics Core using a Fusion Orbitrap Mass Spectrometer.

### PfGRP170-BirA biotinylation and mass spectrometry

To confirm that proteins were biotinylated when biotin was added to the PfGRP170-BirA parasites, parasites were incubated 24 hours in media containing 2.5nM WR + 150μg of biotin (Sigma). Parasites were isolated using 0.04% saponin in 1X PBS and the lysates were analyzed via western blot as described above. Secondary antibodies used were IRDye 680CW goat anti-rabbit IgG and IRDye 800CW Streptavidin (LICOR). 3D7 parasites incubated with media containing 150μg of biotin for 24 hours was used as a control.

For PfGRP170-HA-BirA streptavidin IP’s, cultures were incubated for 24 hours in media containing 2.5nM WR + 150μg of biotin (Sigma). 48 mL of asynchronous culture at high parasitemia (10% or higher) were harvested for IP as described above with the following modifications. Streptavidin MagneSphere Paramagnetic Particle beads (Promega) were used to isolated biotinylated proteins. To prepare the Streptavidin beads for IP, beads were washed three times in 1 mL of 1X PBS. Incubations of lysate with the magnetic beads were performed at room temperature for 30 minutes. After the unbound fraction was removed, beads were washed twice in 8M Urea (150mM NaCL, 50mM Tris HCL pH 7.4) and once in 1X PBS. The biotinylated proteins on magnetic beads were digested with trypsin and analyzed at the Emory University Integrated Proteomics Core using a Fusion Orbitrap Mass Spectrometer. 3D7 control streptavidin IP’s were conducted as above but without the addition of 2.5nM WR to the media.

### Proximity Ligation Assays

Asynchronous PfGRP170-GFP-DDD parasites were fixed as described above, approximately 24 hours after the removal of TMP. The proximity ligation assay was performed using the Duolink PLA Fluorescence kit (Sigma) per the manufacturers protocol. For the BiP/PfGRP170 PLA assay, primary antibodies mouse anti-GFP (Roche 11814460001, 1:500) and rabbit anti-BiP MRA-1246 (BEI resources (1:100) were used. For the negative control primary antibodies mouse anti-plasmepsin V (From D. Goldberg, 1:1) and rabbit anti-GFP A-6455 (Invitrogen, 1:200) were used.

### Ring Stage Survival Assay

The ring-stage survival assay method was performed on 3D7 (control) and PfGRP170-BirA parasites as described previously, with a slight adjustment^80^. Cultures were synchronized using 5% sorbitol (Sigma-Aldrich, St. Louis, MO, USA), pre-warmed to 37°C, to obtain the highest proportion of rings, ≥50%. The cultures were placed back under previously described conditions for 24 hours and followed-up the next morning. Thin blood smears were methanol fixed and stained with 10% Giemsa for 15 minutes and evaluated for mature schizonts with visible nuclei (10-12). The parasites were independently suspended in PRMI-1640 supplemented with 15U/ml of sodium heparin (Sigma-Aldrich, St. Louis, MO, USA) to disrupt spontaneous rosettes formation for 15 minutes at 37 °C. After incubation, each parasite culture was layered onto a 75/25% percoll (GE Healthcare Life Sciences, Pittsburgh, PA, USA) gradient, and centrifuged at 3000rpm for 15 minutes. The intermediate phases containing the mature schizonts of each culture, were independently collected, gently washed in RPMI and transferred into two new T25 flasks with fresh cRPMI and erythrocytes for 3-hour incubation at previously described conditions. Thin blood smears were prepared as previously described, to ensure >10% schizonts count.

At the 3-hour mark, the parasites were taken-out of incubation and treated with 5% sorbitol to remove the remaining mature schizonts, which had not invaded erythrocytes yet. Parasitemia was adjusted to 1% at 2% hematocrit by adding uninfected erythrocytes and cPRMI, after the evaluation of quick stained Giemsa smears. The parasites were exposed to 700nM DHA or 1% dimethyl sulfoxide (DMSO) for 6 hours. After the 6-hour incubation period, the parasites were washed to remove the drug or DMSO and re-suspended in 1ml of cRPMI. The parasites were then transferred into two new well in the 48-well culture plate, incubated at 37 °C under a 90 % N2, 5 % CO2, and 5 % O2 gas mixture for 66 hours, after which thin blood smears were prepared, methanol fixed, stained with 10% Giemsa for 15 minutes and read by three operators. Growth rate and percent survival was calculated by counting the number of parasitized cells in an estimated 2000 erythrocytes.

## REFERENCES

1 World Health Organization. World Malaria Report. (2017).

2 Amaratunga, C., Witkowski, B., Khim, N., Menard, D. & Fairhurst, R. M. Artemisinin resistance in Plasmodium falciparum. The Lancet Infectious Disease 14, 449–450 (2014).

3 Dondorp, A. M. et al. Artemisinin resistance in Plasmodium falciparum malaria. N Engl J Med 361, 455–467, doi:10.1056/NEJMoa0808859 (2009).

4 Roper, C. et al. Intercontinental spread of pyrimethamine-resistant malaria. Science 305, 1124 (2004).

5 Wootton, J. C. et al. Genetic diversity and chloroquine selective sweeps in Plasmodium falciparum. Nature 418, 320–323 (2002).

6 Mita, T. et al. Limited geographical origin and global spread of sulfadoxine-resistant dhps alleles in Plasmodium falciparum populations. J Infect Dis 204, 1980–1988, doi:10.1093/infdis/jir664 (2011).

7 Miller, L. H., Baruch, D. I., Marsh, K. & Doumbo, O. K. The pathogenic basis of malaria. Nature 415, 673–679 (2002).

8 Mok, S. et al. Drug resistance. Population transcriptomics of human malaria parasites reveals the mechanism of artemisinin resistance. Science 347, 431–435, doi:10.1126/science.1260403 (2015).

9 Rocamora, F. et al. Oxidative stress and protein damage responses mediate artemisinin resistance in malaria parasites. PLoS Pathog 14, e1006930, doi:10.1371/journal.ppat.1006930 (2018).

10 Zhang, M. et al. Inhibiting the Plasmodium eIF2alpha Kinase PK4 Prevents Artemisinin-Induced Latency. Cell Host Microbe 22, 766–776 e764, doi:10.1016/j.chom.2017.11.005 (2017).

11 Deponte, M. et al. Wherever I may roam: protein and membrane trafficking in P. falciparum-infected red blood cells. Mol Biochem Parasitol 186, 95–116, doi:10.1016/j.molbiopara.2012.09.007 (2012).

12 Boddey, J. A. & Cowman, A. F. Plasmodium nesting: remaking the erythrocyte from the inside out. Annu Rev Microbiol 67, 243–269, doi:10.1146/annurev-micro-092412-155730 (2013).

13 Boddey, J. A. et al. Role of plasmepsin V in export of diverse protein families from the Plasmodium falciparum exportome. Traffic 14, 532–550, doi:10.1111/tra.12053 (2013).

14 Russo, I. et al. Plasmepsin V licenses Plasmodium proteins for export into the host erythrocyte. Nature 463, 632–636, doi:10.1038/nature08726 (2010).

15 Tonkin, C. J., Kalanon, M. & McFadden, G. I. Protein targeting to the malaria parasite plastid. Traffic 9, 166–175, doi:10.1111/j.1600-0854.2007.00660.x (2008).

16 Zhang, M. et al. PK4, a eukaryotic initiation factor 2α(eIF2α) kinase, is essential for the development of the erythrocytic cycle of Plasmodium. PNAS 109, 3956–3961 (2012).

17 Walter, P. & David, R. The unfolded protein response: from stress pathway to homeostatic regulation. Science 334, 1081–1086 (2011).

18 Araki, K. & Nagata, K. Protein folding and quality control in the ER. Cold Spring Harb Perspect Biol 3, a007526, doi:10.1101/cshperspect.a007526 (2011).

19 Gidalevitz, T., Stevens, F. & Argon, Y. Orchestration of secretory protein folding by ER chaperones. Biochim Biophys Acta 1833, 2410–2424, doi:10.1016/j.bbamcr.2013.03.007 (2013).

20 Hotamisligil, G. S. Endoplasmic reticulum stress and atherosclerosis. Nat Med 16, 396–399, doi:10.1038/nm0410-396 (2010).

21 Xu, C., Bailly-Maitre, B. & Reed, J. C. Endoplasmic reticulum stress: cell life and death decisions. J Clin Invest 115, 2656–2664, doi:10.1172/JCI26373 (2005).

22 Rutkowski, D. T. & Hegde, R. S. Regulation of basal cellular physiology by the homeostatic unfolded protein response. J Cell Biol 189, 783–794, doi:10.1083/jcb.201003138 (2010).

23 Wang, H. et al. The Endoplasmic Reticulum Chaperone GRP170: From Immunobiology to Cancer Therapeutics. Front Oncol 4, 377, doi:10.3389/fonc.2014.00377 (2014).

24 Pavithra, S. R., Kumar, R. & Tatu, U. Systems analysis of chaperone networks in the malarial parasite Plasmodium falciparum. PLoS Comput Biol 3, 1701–1715, doi:10.1371/journal.pcbi.0030168 (2007).

25 Shonhai, A., Boshoff, A. & Blatch, G. L. The structural and functional diversity of Hsp70 proteins from Plasmodium falciparum. Protein Sci 16, 1803–1818, doi:10.1110/ps.072918107 (2007).

26 Andreasson, C., Rampelt, H., Fiaux, J., Druffel-Augustin, S. & Bukau, B. The endoplasmic reticulum Grp170 acts as a nucleotide exchange factor of Hsp70 via a mechanism similar to that of the cytosolic Hsp110. J Biol Chem 285, 12445–12453, doi:10.1074/jbc.M109.096735 (2010).

27 de Keyzer, J., Steel, G. J., Hale, S. J., Humphries, D. & Stirling, C. J. Nucleotide binding by Lhs1p is essential for its nucleotide exchange activity and for function in vivo. J Biol Chem 284, 31564–31571, doi:10.1074/jbc.M109.055160 (2009).

28 Behnke, J. & Hendershot, L. M. The large Hsp70 Grp170 binds to unfolded protein substrates in vivo with a regulation distinct from conventional Hsp70s. J Biol Chem 289, 2899–2907, doi:10.1074/jbc.M113.507491 (2014).

29 Park, J. et al. The Chaperoning Properties of Mouse Grp170, a Member of the Third Family of Hsp70 Related Proteins. Biochemistry 42, 14893–14902 (2003).

30 Buck, T. M. et al. The Lhs1/GRP170 chaperones facilitate the endoplasmic reticulum-associated degradation of the epithelial sodium channel. J Biol Chem 288, 18366–18380, doi:10.1074/jbc.M113.469882 (2013).

31 Beck, J. R., Muralidharan, V., Oksman, A. & Goldberg, D. E. PTEX component HSP101 mediates export of diverse malaria effectors into host erythrocytes. Nature 511, 592–595, doi:10.1038/nature13574 (2014).

32 Florentin, A. et al. PfClpC Is an Essential Clp Chaperone Required for Plastid Integrity and Clp Protease Stability in Plasmodium falciparum. Cell Rep 21, 1746–1756, doi:10.1016/j.celrep.2017.10.081 (2017).

33 Muralidharan, V., Oksman, A., Pal, P., Lindquist, S. & Goldberg, D. E. Plasmodium falciparum heat shock protein 110 stabilizes the asparagine repeat-rich parasite proteome during malarial fevers. Nat Commun 3, 1310–1310 (2012).

34 Easton, D. P., Kaneko, Y. & Subjeck, J. R. The hsp110 and Grp170 stress proteins: newly recognized relatives of the Hsp70s. Cell Stress Chaperones 5, 276–290 (2000).

35 Cobb, D. W. et al. The Exported Chaperone PfHsp70x Is Dispensable for the Plasmodium falciparum Intraerythrocytic Life Cycle. mSphere 2, doi:10.1128/mSphere.00363-17 (2017).

36 Muralidharan, V., Oksman, A., Iwamoto, M., Wandless, T. J. & Goldberg, D. E. Asparagine repeat function in a Plasmodium falciparum protein assessed via a regulatable fluorescent affinity tag. Proc Natl Acad Sci U S A 108, 4411–4416, doi:10.1073/pnas.1018449108 (2011).

37 Ganter, M. et al. Plasmodium falciparum CRK4 directs continuous rounds of DNA replication during schizogony. Nat Microbiol 2, 17017, doi:10.1038/nmicrobiol.2017.17 (2017).

38 Ito, D., Schureck, M. A. & Desai, S. A. An essential dual-function complex mediates erythrocyte invasion and channel-mediated nutrient uptake in malaria parasites. Elife 6, doi:10.7554/eLife.23485 (2017).

39 Waller, R. F., Reed, M. B., Cowman, A. F. & McFadden, G. I. Protein trafficking to the plastid of Plasmodium falciparum is via the secretory pathway. EMBO J 19, 1794–1802 (2000).

40 Tonkin, C. J., Struck, N. S., Mullin, K. A., Stimmler, L. M. & McFadden, G. I. Evidence for Golgi-independent transport from the early secretory pathway to the plastid in malaria parasites. Mol Microbiol 61, 614–630, doi:10.1111/j.1365-2958.2006.05244.x (2006).

41 Heiny, S. R., Pautz, S., Recker, M. & Przyborski, J. M. Protein Traffic to the Plasmodium falciparum apicoplast: evidence for a sorting branch point at the Golgi. Traffic 15, 1290–1304, doi:10.1111/tra.12226 (2014).

42 Fellows, J. D., Cipriano, M. J., Agrawal, S. & Striepen, B. A Plastid Protein That Evolved from Ubiquitin and Is Required for Apicoplast Protein Import in Toxoplasma gondii. mBio 8, 1–18 (2017).

43 Sheiner, L. et al. A systematic screen to discover and analyze apicoplast proteins identifies a conserved and essential protein import factor. PLoS Pathog 7, e1002392, doi:10.1371/journal.ppat.1002392 (2011).

44 Yeh, E. & DeRisi, J. L. Chemical rescue of malaria parasites lacking an apicoplast defines organelle function in blood-stage Plasmodium falciparum. PLoS Biol 9, e1001138, doi:10.1371/journal.pbio.1001138 (2011).

45 Chen, A. L. et al. Novel components of the Toxoplasma inner membrane complex revealed by BioID. MBio 6, e02357–02314, doi:10.1128/mBio.02357-14 (2015).

46 Painter, H. J. et al. Genome-wide real-time in vivo transcriptional dynamics during Plasmodium falciparum blood-stage development. Nat Commun 9, 2656, doi:10.1038/s41467-018-04966-3 (2018).

47 Soderberg, O. et al. Direct observation of individual endogenous protein complexes in situ by proximity ligation. Nat Methods 3, 995–1000, doi:10.1038/nmeth947 (2006).

48 Gullberg, M. et al. Cytokine detection by antibody-based proximity ligation. Proc Natl Acad Sci U S A 101, 8420–8424, doi:10.1073/pnas.0400552101 (2004).

49 Fredriksson, S. et al. Protein detection using proximity-dependent DNA ligation assays. Nature Biotechnology 20, 473–477 (2002).

50 Kulzer, S. et al. Plasmodium falciparum-encoded exported hsp70/hsp40 chaperone/co-chaperone complexes within the host erythrocyte. Cell Microbiol 14, 1784–1795, doi:10.1111/j.1462-5822.2012.01840.x (2012).

51 Tyson, J. R. & Stirling, C. J. LHS1 and SIL1 provide a lumenal function that is essential for protein translocation into the endoplasmic reticulum. The EMBO Journal 19, 6440–6452 (2000).

52 Harbut, M. B. et al. Targeting the ERAD pathway via inhibition of signal peptide peptidase for antiparasitic therapeutic design. Proc Natl Acad Sci U S A 109, 21486–21491 (2012).

53 Chaubey, S., Grover, M. & Tatu, U. Endoplasmic reticulum stress triggers gametocytogenesis in the malaria parasite. J Biol Chem 289, 16662–16674, doi:10.1074/jbc.M114.551549 (2014).

54 Fennell, C. et al. PfeIK1, a eukaryotic initiation factor 2alpha kinase of the human malaria parasite Plasmodium falciparum, regulates stress-response to amino-acid starvation. Malar J 8, 99, doi:10.1186/1475-2875-8-99 (2009).

55 Babbitt, S. E. et al. Plasmodium falciparum responds to amino acid starvation by entering into a hibernatory state. Proc Natl Acad Sci U S A 109, E3278–3287 (2012).

56 Ward, P., Equinet, L., Packer, J. & Doerig, C. Protein kinases of the human malaria parasite Plasmodium falciparum: the kinome of a divergent eukaryote. BMC Genomics 5, 79, doi:10.1186/1471-2164-5-79 (2004).

57 Foth, B. J. et al. Dissecting Apicoplast Targeting in the Malaria Parasite Plasmodium falciparum. Science 299, 705–708 (2003).

58 Ramya, T. N., Karmodiya, K., Surolia, A. & Surolia, N. 15-deoxyspergualin primarily targets the trafficking of apicoplast proteins in Plasmodium falciparum. J Biol Chem 282, 6388–6397, doi:10.1074/jbc.M610251200 (2007).

59 Ramya, T. N., Surolia, N. & Surolia, A. 15-Deoxyspergualin modulates Plasmodium falciparum heat shock protein function. Biochem Biophys Res Commun 348, 585–592, doi:10.1016/j.bbrc.2006.07.082 (2006).

60 Gruring, C. et al. Uncovering common principles in protein export of malaria parasites. Cell Host Microbe 12, 717–729, doi:10.1016/j.chom.2012.09.010 (2012).

61 Boddey, J. A. et al. Export of malaria proteins requires co-translational processing of the PEXEL motif independent of phosphatidylinositol-3-phosphate binding. Nat Commun 7, 10470, doi:10.1038/ncomms10470 (2016).

62 Marti, M., Good, R. T., Rug, M., Knuepfer, E. & Cowman, A. F. Targeting malaria virulence and remodeling proteins to the host erythrocyte. Science 306, 1930–1933 (2004).

63 Hiller, N. L. et al. A Host-Targeting Signal in Virulence Proteins Reveals a Secretome in Malarial Infection. Science 306 (2015).

64 Heiny, S. R., Spork, S. & Przyborski, J. The apicoplast of the human malaria parasite P. falciparum. Journal of Endocytobiosis and Cell Research 23, 91–95 (2012).

65 Batinovic, S. et al. An exported protein-interacting complex involved in the trafficking of virulence determinants in Plasmodium-infected erythrocytes. Nat Commun 8, 16044, doi:10.1038/ncomms16044 (2017).

66 Wang, Y. et al. Involvement of oxygen-regulated protein 150 in AMP-activated protein kinase-mediated alleviation of lipid-induced endoplasmic reticulum stress. J Biol Chem 286, 11119–11131, doi:10.1074/jbc.M110.203323 (2011).

67 Sanson, M. et al. Oxidized low-density lipoproteins trigger endoplasmic reticulum stress in vascular cells: prevention by oxygen-regulated protein 150 expression. Circ Res 104, 328–336, doi:10.1161/CIRCRESAHA.108.183749 (2009).

68 Yoshida, H., Matsui, T., Yamamoto, A., Okada, T. & Mori, K. XbP1 mRNA Is Induced by ATF6 and Spliced by IRE1 in Response to ER Stress to Produce a Highly Active Transcription Factor. Cell 107, 881–891, doi:https://doi.org/10.1016/S0092-8674(01)00611-0 (2001).

69 Collins, C. R., Hackett, F., Atid, J., Tan, M. S. Y. & Blackman, M. J. The Plasmodium falciparum pseudoprotease SERA5 regulates the kinetics and efficiency of malaria parasite egress from host erythrocytes. PLoS Pathog 13, e1006453, doi:10.1371/journal.ppat.1006453 (2017).

70 Ruecker, A. et al. Proteolytic activation of the essential parasitophorous vacuole cysteine protease SERA6 accompanies malaria parasite egress from its host erythrocyte. J Biol Chem 287, 37949–37963, doi:10.1074/jbc.M112.400820 (2012).

71 Thomas, J. A. et al. A protease cascade regulates release of the human malaria parasite Plasmodium falciparum from host red blood cells. Nat Microbiol 3, 447–455, doi:10.1038/s41564-018-0111-0 (2018).

72 Zhao, X. et al. PfRON3 is an erythrocyte-binding protein and a potential blood-stage vaccine candidate antigen. Malaria Journal 13, 490, doi:10.1186/1475-2875-13-490 (2014).

73 Ling, I. T. et al. The Plasmodium falciparum clag9 gene encodes a rhoptry protein that is transferred to the host erythrocyte upon invasion. Mol Microbiol 52, 107–118, doi:10.1111/j.1365-2958.2003.03969.x (2004).

74 Trenholme, K. R. et al. clag9: A cytoadherence gene in Plasmodium falciparum essential for binding of parasitized erythrocytes to CD36. Proceedings of the National Academy of Sciences of the United States of America 97, 4029–4033, doi:10.1073/pnas.040561197 (2000).

75 Balu, B., Shoue, D. A., Fraser, M. J., Jr. & Adams, J. H. High-efficiency transformation of Plasmodium falciparum by the lepidopteran transposable element piggyBac. Proc Natl Acad Sci U S A 102, 16391–16396, doi:10.1073/pnas.0504679102 (2005).

76 van Ooij, C. et al. The malaria secretome: from algorithms to essential function in blood stage infection. PLoS Pathog 4, e1000084, doi:10.1371/journal.ppat.1000084 (2008).

77 Li, M. Z. & Elledge, S. J. Harnessing homologous recombination in vitro to generate recombinant DNA via SLIC. Nat Methods 4, 251–256, doi:10.1038/nmeth1010 (2007).

78 Iwanaga, S., Kato, T., Kaneko, I. & Yuda, M. Centromere plasmid: a new genetic tool for the study of Plasmodium falciparum. PLoS One 7, e33326, doi:10.1371/journal.pone.0033326 (2012).

79 Ganesan, S. M. et al. Yeast dihydroorotate dehydrogenase as a new selectable marker for Plasmodium falciparum transfection. Mol Biochem Parasitol 177, 29–34, doi:10.1016/j.molbiopara.2011.01.004 (2011).

80 Witkowski, B. et al. Novel phenotypic assays for the detection of artemisinin-resistant Plasmodium falciparum malaria in Cambodia: in-vitro and ex-vivo drug-response studies. The Lancet Infectious Diseases 13, 1043–1049, doi:10.1016/s1473-3099(13)70252-4 (2013).

